# The giant Mimivirus 1.2 Mb genome is elegantly organized into a 30 nm helical protein shield

**DOI:** 10.1101/2022.02.17.480895

**Authors:** Alejandro Villalta, Alain Schmitt, Leandro F. Estrozi, Emmanuelle R. J. Quemin, Jean-Marie Alempic, Audrey Lartigue, Vojtěch Pražák, Lucid Belmudes, Daven Vasishtan, Agathe M. G. Colmant, Flora A. Honoré, Yohann Couté, Kay Grünewald, Chantal Abergel

**Author notes:** These authors contributed equally to the work (Alejandro Villalta, Alain Schmitt, Leandro F. Estrozi). Department of Virology, Institute for Integrative Biology of the Cell (I2BC), Centre National de la Recherche Scientifique UMR9198, 1, avenue de la Terrasse, 91198 Gif-sur-Yvette Cedex, France (Emmanuelle R. J. Quemin).

## Abstract

Mimivirus is the prototype of the *Mimiviridae* family of giant dsDNA viruses. Little is known about the organization of the 1.2 Mb genome inside the membrane-limited nucleoid filling the ∼0.5 µm icosahedral capsids. Cryo-electron microscopy, cryo-electron tomography and proteomics revealed that it is encased into a ∼30 nm diameter helical protein shell surprisingly composed of two GMC-type oxidoreductases, which also form the glycosylated fibrils decorating the capsid. The genome is arranged in 5- or 6-start left-handed super-helices, with each DNA-strand lining the central channel. This luminal channel of the nucleoprotein fiber is wide enough to accommodate oxidative stress proteins and RNA polymerase subunits identified by proteomics. Such elegant supramolecular organization would represent a remarkable evolutionary strategy for packaging and protecting the genome, in a state ready for immediate transcription upon unwinding in the host cytoplasm. The parsimonious use of the same protein in two unrelated substructures of the virion is unexpected for a giant virus with thousand genes at its disposal.

**One-Sentence Summary:** Mimivirus genome organization in the icosahedral virion.

## Introduction

Giant viruses were discovered with the isolation of Mimivirus from Acanthamoeba species (*1, 2*). These viruses now represent a highly diverse group of dsDNA viruses infecting unicellular eukaryotes (*3*) which play important roles in the environment (*4–6*). They also challenge the canonical definitions of viruses (*7, 8*) as they can encode central translation components (*2, 9*) as well as a complete glycosylation machinery (*10, 11*) among other unique features.

Mimivirus has been the most extensively studied giant virus over the years (*12*). The virions are 0.75 µm wide and consist of icosahedral capsids of 0.45 µm diameter surrounded by a dense layer of radially arranged fibrils (*2*). Structural analyses of the virions have provided some insights into the capsid structure (*13–17*) but given the size of the icosahedral particles (and hence the sample thickness), accessing the internal organization of the core of the virions remains challenging. Consequently, little is known about the packaging of the 1.2 Mb dsDNA genome (*18*). Inside the capsids, a lipid membrane delineates an internal compartment (∼340 nm in diameter, Fig 1A, Fig. S1A), referred to as the nucleoid, which contains the viral genome, together with all proteins necessary to initiate the replicative cycle within the host cytoplasm (*17, 19–21*).

**Figure 1:**
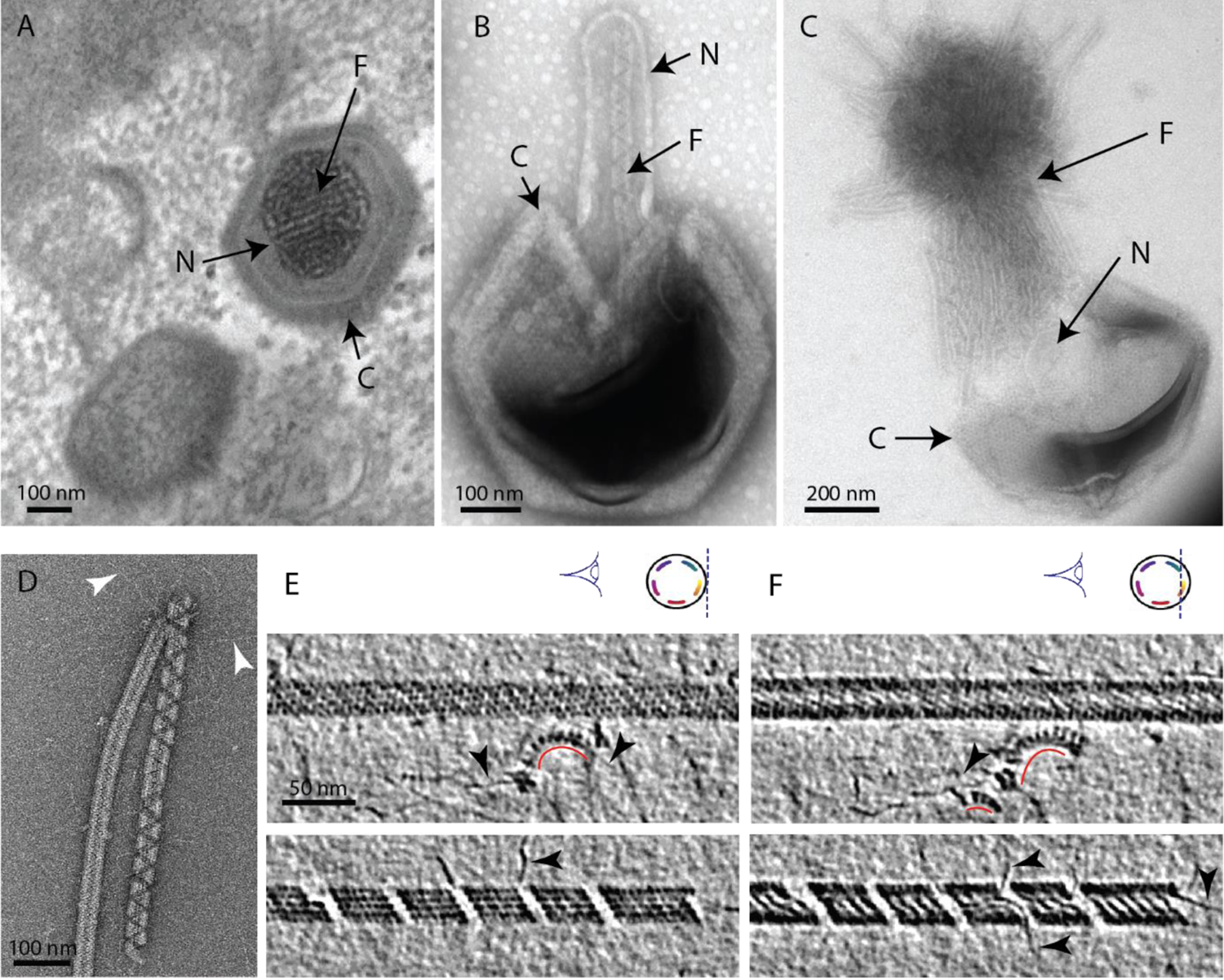
The Mimivirus genomic fiber. A] Micrograph of an ultrathin section of resin-embedded infected cells showing the DNA tightly packed inside Mimivirus capsids (C) with electron dense material inside the nucleoid (N). The string-like features, most likely enhanced by the dehydration caused by the fixation and embedding protocol, correspond to the genomic fiber (F) packed into the nucleoid. B] Micrograph of negative stained Mimivirus capsid (C) opened *in vitro* with the genomic fiber (F) still being encased into the membrane limited nucleoid (N). C] Multiple strands of the flexible genomic fiber (F) are released from the capsid (C) upon proteolytic treatment. D] Micrograph of negative stained purified Mimivirus genomic fibers showing two conformations (the right fiber resembling the one in [B] and free DNA strands (white arrowheads). E] Slices through two electron cryo tomograms of the isolated helical protein shell of the purified genomic fibers in compact (top) or relaxed conformation (bottom) in the process of losing one protein strand (Fig. S11, Tomo. S1 & S4). F] Different slices through the two tomograms shown in [E] reveal DNA strands lining the helical protein shell of the purified genomic fibers in compact (top) or relaxed conformation (bottom). Examples of DNA strands extending out at the breaking points of the genomic fiber are marked by black arrowheads. Note in the top panel, individual DNA strands coated by proteins (red arc). The slicing planes at which the Mimivirus genomic fibers were viewed are indicated on diagrams on the top right corner as blue dashed lines and the internal colored segments correspond to DNA strands lining the protein shell. The thickness of the tomographic slices is 1.1 nm and the distance between tomographic slices in panels [E] and [F] is 4.4 nm. Scale bars as indicated.

Acanthamoeba cells engulf Mimivirus particles, fooled by their bacteria-like size and the heavily glycosylated decorating fibrils (*1, 2, 11*). Once in the phagosome, the Stargate portal located at one specific vertex of the icosahedron opens up (*22*), enabling the viral membrane to fuse with that of the host vacuole to deliver the nucleoid into the host cytoplasm (*17, 20*). EM studies have shown that next the nucleoid gradually loses its electron dense appearance, transcription begins and the early viral factory is formed (*21, 23, 24*). Previous Atomic Force Microscopy (AFM) studies of the Mimivirus infectious cycle suggested that the DNA forms a highly condensed nucleoprotein complex enclosed within the nucleoid (*13*). Here we show that opening of the large icosahedral capsid *in vitro* led to the release of rod-shaped structures of about 30 nm width. These structures were further purified and the various conformations characterized using cryo-electron microscopy (cryo-EM), tomography and MS-based proteomics.

## Results

### Capsid opening induces the release of a ∼ 30 nm-wide rod-shaped structure that contains the dsDNA genome

We developed an *in vitro* protocol for particle opening that led to the release of ∼ 30 nm-wide rod-shaped fibers of several microns in length (Fig. 1, Fig. S1). We coined this structure the Mimivirus genomic fiber. Complete expulsion of the opened nucleoid content resulted in the observation of bundled fibers resembling a “ball of yarn” (*13*) (Fig. 1C,). The used capsid opening procedure involves limited proteolysis and avoids harsh conditions, as we found that the structure becomes completely denatured by heat (95°C) and is also sensitive to acidic treatment, thus preventing its detection in such conditions (*17*). Various conformations of the genomic fiber were observed, sometimes even on a same fiber (Fig. 1D), ranging from the most compact rod-shaped structures (Fig. 1D (left), E-F (top)) to more relaxed structures where DNA strands begin to dissociate (Fig. 1D (right), E-F (bottom)). After optimizing the *in vitro* extraction on different strains of group-A Mimiviruses, we focused on an isolate from La Réunion Island (Mimivirus reunion), as more capsids were opened by our protocol, leading to higher yields of genomic fibers that were subsequently purified on sucrose gradient. All opened capsids released genomic fibers (Fig. 1C, Fig. S1B).

The first confirmation of the presence of DNA in the genomic fiber was obtained by agarose gel electrophoresis (Fig. S2). Electron cryo microscopy (cryo-EM) bubblegram analysis (*25, 26*) gave a further indication that the nucleic acid is located in the fiber lumen. Alike other nucleoprotein complexes, fibers are expected to be more susceptible to radiation damage then pure proteinaceous structures. Surprisingly, the specimen could sustain higher electron irradiation before the appearance of bubbles compared to other studies (*27*): 600 e-/Å² for relaxed helices and up to 900 e-/Å² for long compact ones, while no bubbles could be detected in unfolded ribbons (Fig. S3). For comparison, bacteriophage capsids containing free DNA, i.e. not in the form of nucleoproteins, show bubbling for doses of ∼30-40 e-/Å² (*27*).

### Cryo-EM single particle analysis of the different compaction states of the Mimivirus genomic fiber

In order to shed light on the Mimivirus genome packaging strategy and to determine the structure of the purified genomic fibers, we performed cryo-EM single-particle analysis. The different conformations of the genomic fiber initially observed by negative staining (Fig. 1) and cryo-EM resulted in a highly heterogeneous dataset for single-particle analysis. In order to separate different conformations, *in silico* sorting through 2D classification (using Relion (*28, 29*)) was performed. Next, we carried out cluster analysis relying on the widths of the helical segments and correlations (real and reciprocal spaces) between the experimental 2D class patterns (Figs. S4 & S5). Three independent clusters (Cl) could be distinguished, corresponding to the compact (Cl1), intermediate (Cl2) and relaxed (Cl3) fiber conformations (Fig. S5) with the latter being the widest. For each cluster, we determined their helical symmetry parameters by image power spectra analyses and performed structure determination and refinement (Fig. 2, Fig. S6-8).

**Figure 2:**
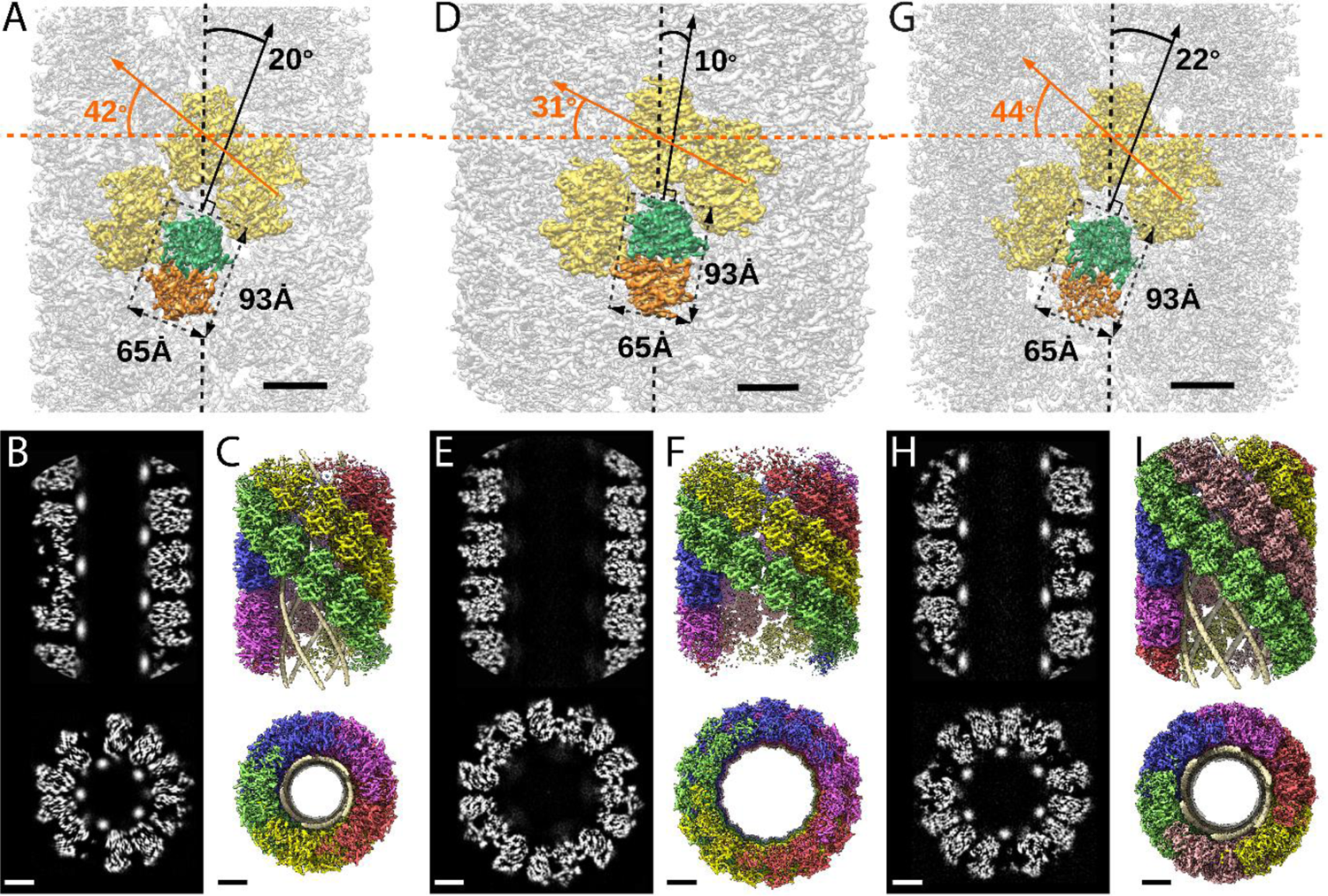
Structures of the Mimivirus genomic fiber for Cl1a [A-C], Cl3a [D-F], and Cl2 [G-I]: A, D, G] Electron microscopy (EM) maps of Cl1a (A) and Cl3a (D) are shown with each monomer of one GMC-oxidoreductase dimer colored in green and orange and 3 adjacent dimers in yellow, to indicate the large conformational change taking place between the two fiber states. The transition from Cl1a to Cl3a (5-start helix) corresponds to a rotation of each individual unit (corresponding to a GMC-oxidoreductase dimer) by ∼ −10° relative to the fiber longitudinal axis and a change in the steepness of the helical rise by ∼ −11°. Scale bars 50 Å. Compared to Cl1a, the Cl2 6-start helix (G) shows a difference of ∼2° relative to the fiber longitudinal axis and ∼2° in the steepness of the helical rise. Scale bars 50 Å. B, E, H] Cross-sectional (bottom) and longitudinal (top) sections through the middle of final Cl1a (B) Cl3a (E) and Cl2 (H) EM maps. Scale bars 50 Å. C, F, I] Longitudinal (top) and orthogonal (bottom) views of final Cl1a (C) and Cl3a (F) and Cl2 (I) EM maps color-coded according to each start of the 5-start helix. Densities for some asymmetric units in the front have been removed on the side view map to show the 5 DNA strands lining the protein shell interior. Scale bars 50 Å.

For both Cl1a and Cl3a conformations, after 3D-refinement, we obtained helical structures of 3.7 Å resolution (FSC threshold 0.5, masked), corresponding to 5-start left-handed helices made of a ∼8 nm-thick proteinaceous external shell (Figs. S6 and S7). For the most compact conformation (Cl1a) 5 dsDNA strands were lining the interior of the protein shell leaving a ∼9 nm wide central channel (Fig. 2-3, Table S1). The DNA strands appear as curved cylinders in the helical structure, the characteristic shape of the DNA (minor and major groove) becoming only visible after focused refinement of a single DNA strand (Fig. 3, Fig. S9-S10). In the relaxed subcluster Cl3a, the DNA strands at the interface to the ∼17 nm-wide central channel are not clearly recognizable (Fig. 2, Table S1 & Fig. S7), most likely because they are at least partially detached inside the broken expanded fiber. The breaks after relaxation of the helix might be the result of the extraction and purification treatment, while DNA will remain in the central channel, at least in the early phase of Acanthamoeba infection.

**Figure 3:**
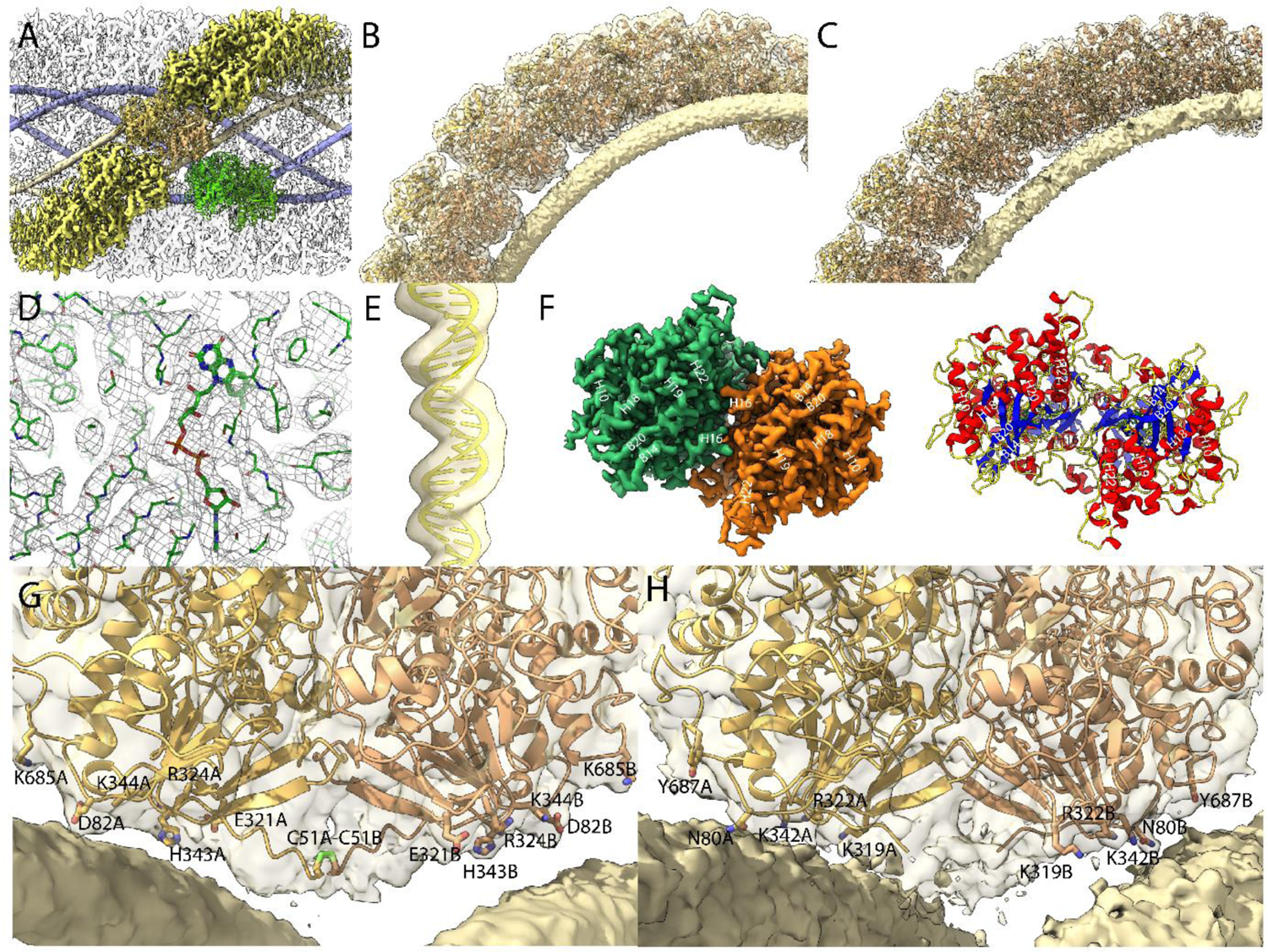
Maps of the compact (Cl1a and Cl2) genomic fiber structures: A] Cl1a EM map corresponding to the protein shell prior focused refinement is shown as a transparent surface and the 5 DNA strands as solid surface. One protein dimer strand is shown yellow except for one asymmetric unit (transparent yellow) to illustrate the dimer fit. The position of a second dimer (green) from the adjacent dimer strand is shown to emphasize that the DNA strand (gold) is lining the interface between two dimers. B, C] Cartoon representation of GMC-oxidoreductase qu_946 dimers fitted into one of the Cl1a 5-start helix strands in the 3.7 Å resolution map (B) and GMC-oxidoreductase. 143 dimers fitted into one of the Cl2 6-start helix strands in the respective 4.0 Å resolution map (C). The maps are shown at a threshold highlighting the periodicity of contacts between the dsDNA and the protein shell. D] Zoom into the 3.3 Å resolution focused refined Cl1a map illustrating the fit of the side chains and the FAD ligand. E] Cartoon representation of the DNA fitted in the focused refined DNA only map (Fig. S10). F] Focused refined Cl1a map colored by monomer, next to a cartoon representation of the qu_946 dimer (α-helices in red, β-strands in blue and coils in yellow). Secondary structure elements are annotated in both representations (H: helix, B: beta-strand). G, H] Cartoon representation of qu_946 (G) and qu_143 (H) fitted into their respective cryo-EM maps highlighting the interacting residues (given as stick models) between each monomer and one dsDNA strand. The isosurface threshold chosen allows visualization of density for the manually built N-terminal residues including the residues C51 (stick model) of two neighboring monomers that form a terminal disulfide bridge.

Finally, the 4 Å resolution Cl2 map obtained after 3D refinement (Fig. S8) corresponds to a 6-start left-handed helix made of a ∼8 nm-thick proteinaceous external shell, with 6 dsDNA strands lining the shell interior and leaving a ∼12 nm wide inner channel (Fig. 2, Table S1).

### The most abundant proteins in the genomic fiber are GMC-oxidoreductases, the same that compose the fibrils decorating Mimivirus capsid

MS-based proteomic analyses performed on three biological replicates identified two GMC-oxidoreductases as the main components of the purified genomic fiber (qu_946 and qu_143 in Mimivirus reunion corresponding to L894/93 and R135 respectively in Mimivirus prototype) (Table S2). The available Mimivirus R135 GMC-oxidoreductase dimeric structure (*30*) (PDB 4Z24, lacking the 50 amino acid long cysteine-rich N-terminal domain) was fitted into the EM maps (Figs. 2-3). This is quite unexpected, since GMC-oxidoreductases are already known to compose the fibrils surrounding Mimivirus capsids (*11, 31*). Notably, the proteomic analyses provided different sequence coverages for the GMC-oxidoreductases depending on whether samples were virions or the purified genomic fiber preparations, with substantial under-representation of the N-terminal domain in the genomic fiber (Fig. S12). Accordingly, the maturation of the GMC-oxidoreductases involved in genome packaging must be mediated by one of the many proteases encoded by the virus.

### Analysis of the genomic fiber structure

The EM maps and FSC curves of Cl1a are shown in Fig. S6. An additional step of refinement focused on the asymmetric unit further improved the local resolution to 3.3 Å as indicated by the corresponding FSC (Figs. 3 and S6). After fitting the most abundant GMC-oxidoreductase qu_946 (SWISS-MODEL model (*32*)) in the final map of Cl1a, five additional N-terminal residues in each monomer were manually built using the uninterpreted density available. This strikingly brings the cysteines of each monomer (C51 in qu_946) close enough to allow a disulfide bridge, directly after the 50 amino acids domain not covered in the proteomic analysis of the genomic fiber (Fig. 3G, Fig. S12).

The N-terminal chain, being more disordered than the rest of the structure, is absent from the focused refined map, suggesting that a break of the disulfide bridge could be involved in the observed unwinding process. Models of the three helical assemblies and asymmetric unit were further refined using the real-space refinement program in PHENIX 1.18.2 (*33*). In the 3.3 Å resolution map of the asymmetric unit, most side chains and notably the FAD co-factor are accommodated by density suggesting that the oxidoreductase enzyme could be active (Fig. 3D). Density that can be attributed to the FAD cofactor is also present in the Cl2 and Cl3a maps. The atomic models of Cl1a and Cl3a dimers are superimposable with a core RMSD of 0.68 Å based on Cα atoms.

Inspection of individual genomic fibers in the tomograms confirmed the co-existence of both 5- and 6-start left-handed helices containing DNA (Fig. S11 & Video S1, Tomo. S1). Further, some intermediate and relaxed structures were also observed in which the DNA segments appeared detached from the protein shell and sometimes completely absent from the central channel of the broken fibers. Both GMC-oxidoreductases (qu_946 and qu_143) can be fitted in the 5- and 6-start maps.

In relaxed or broken fibers, large electron dense structures that might correspond to proteins inside the lumen were sometimes visible (Fig. S11C & E & Tomo. S3) as well as dissociating DNA fragments, either in the central channel (Fig. S11B & Tomo. S2) or at the breakage points of the fibers in its periphery (Fig. S11D & Tomo. S4). Densities corresponding to the dimer subunits composing the protein shell were also commonly observed on dissociated DNA strands (Fig. 1E-F, Fig. S11 and Tomo. S1-S4).

In the Cl1a and Cl2 helices, the monomers in each dimer are interacting with two different dsDNA strands. As a result, the DNA strands are interspersed between two dimers, each also corresponding to a different strand of the protein shell helix (Fig. 3). Based on the periodic contacts between protein shell and DNA strands, these interactions might involve, in the case of the Cl1a helix, one aspartate (D82 relative to the N-terminal Methionine in qu_946), one glutamate (E321), three lysines (K84, K344, K685), one arginine (R324) and a histidine (H343) or one asparagine (N80), two lysines (K319, K342), one arginine (R322) and one tyrosine (Y687), in the case of the Cl2 helix (Fig. 3G,H).

Despite the conformational heterogeneity and the flexibility of the rod-shaped structure, we were able to build three atomic models of the Mimivirus genomic fiber, in compact (5- and 6-start) and relaxed (5-start) states. Higher resolution data would still needed to determine the precise structure of the dsDNA corresponding to the viral genome (Fig. 3B, C, E), however, the lower resolution for this part of the map even in focused refinement runs (Fig. 3E) might also mean that the DNA does not always bind in the same orientation.

### Rough estimation of genome compaction to fit into the nucleoid

Since there is a mixture of 5 and 6 strands of DNA in the genomic fiber, this could correspond to 5 or 6 genomes per fiber or to a single folded genome. Assuming that the length of DNA in B-form is ∼34 Å for 10 bp, the Mimivirus linear genome of 1.2×10^6^ bp would extend over ∼400 µm and occupy a volume of 1.3×10^6^ nm^3^ (∼300 µm and ∼1×10^6^ nm^3^ if in A-form) (*34*). The volume of the nucleoid (*19*) (∼ 340 nm in diameter) is approximately 2.1×10^7^ nm^3^ and could accommodate over 12 copies of viral genomes in a naked state, but only 40 µm of the ∼30 nm wide flexible genomic fiber. Obviously, the Mimivirus genome cannot be simply arranged linearly in the genomic fiber and must undergo further compaction to accommodate the 1.2 Mb genome in a ∼40 µm long genomic fiber. As a result, the Mimivirus genome is probably organized as an helical shell internally lined by folded DNA strands, surprisingly resembling the archaeal APBV1 virion structure (*35*).

### Additional proteins, including RNA polymerase subunits, are enriched in the genomic fiber

The proteomic analysis of fiber preparations revealed the presence of additional proteins including several RNA polymerase subunits: Rpb1 and Rpb2 (qu_530/532 and qu_261/259/257/255), Rpb3/11 (qu_493), Rpb5 (qu_245), RpbN (qu_379) and Rpb9 (qu_219), in addition to a kinesin (qu_313), a regulator of chromosome condensation (qu_366), a helicase (qu_572), to be possibly associated with the genome (Table S2). In addition to the two GMC-oxidoreductases, at least three oxidative stress proteins were also identified together with hypothetical proteins (Table S2). As expected, the core protein (qu_431) composing the nucleoid and the major capsid proteins (MCP, qu_446) were significantly decreased in the genomic fiber proteome compared to intact virions. In fact, qu_431 and qu_446 represent respectively 4.5% and 9.4% of the total protein abundance in virions whereas they only account for 0.4% and 0.7% of the total protein abundance in the genomic fiber, suggesting that they could be contaminants in this preparation. On the contrary, we calculated enrichment factors of more than five hundred (qu_946) and twenty-six (qu_143) in the genomic fiber samples compared to the intact virion. Finally, the most abundant RNA polymerase subunit (qu_245) is increased by a factor of eight in the genomic fiber compared to intact virion (if the six different subunits identified are used, the global enrichment is seven-fold). Furthermore, upon inspection of the negative staining micrographs, macromolecules strikingly resembling the characteristic structure of the poxviruses RNA polymerase (*36*) were frequently observed scattered around the unwinding fiber and sometimes sitting on DNA strands near broken fibers (Fig. 4). Together with the tomograms showing large electron dense structures in the lumen, some RNA polymerase units could occupy the center of the genomic fiber.

**Figure 4:**
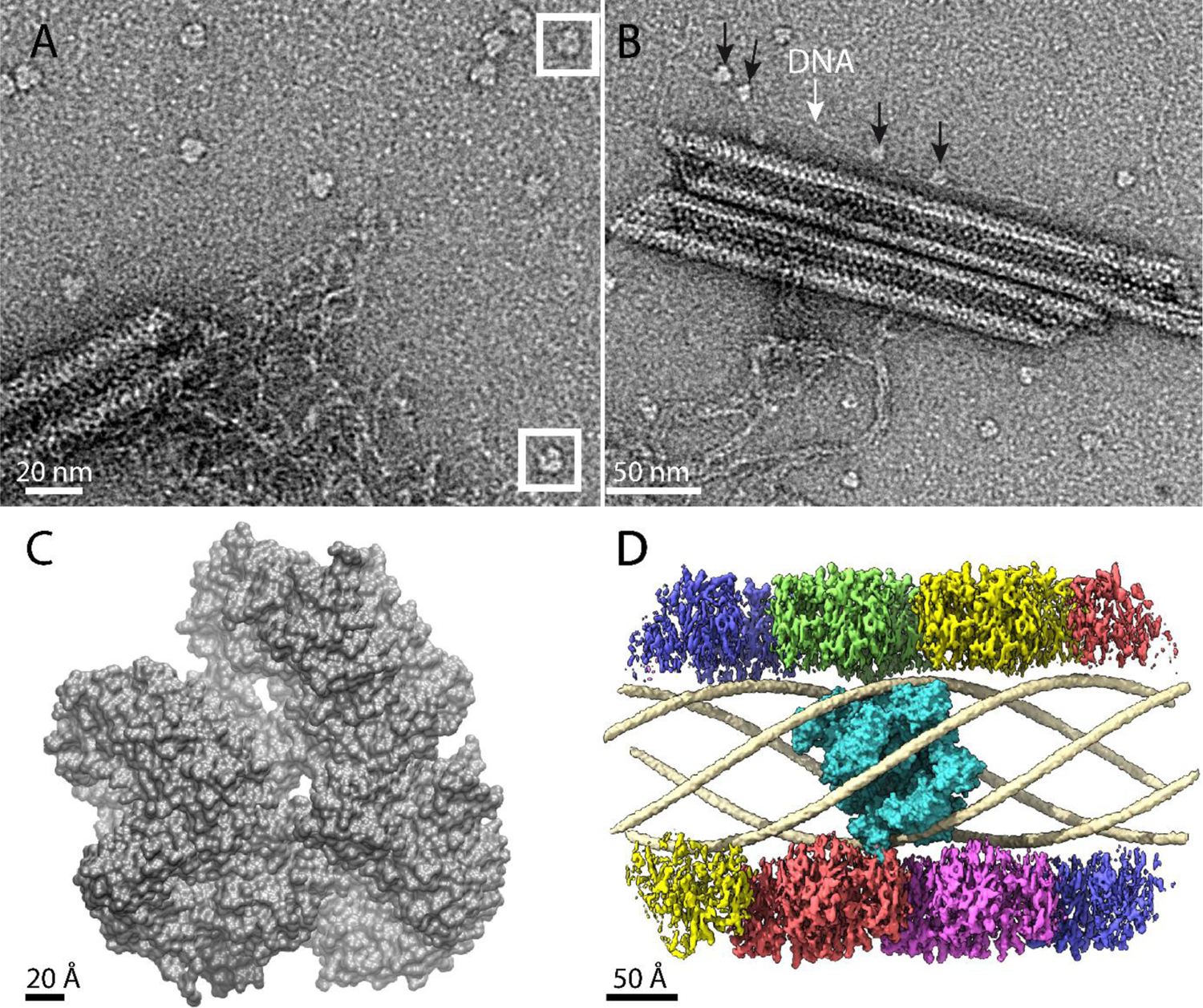
RNA polymerase could be associated to the genomic fiber. A] Micrograph of negative stained fiber with released DNA still being connected to a relaxed and broken fiber and adjacent scattered macromolecular complexes that might resemble RNA polymerases. B] Strikingly, some of them (black arrows) appear to sit on a DNA strand (white arrow). C] Surface representation of the vaccinia virus RNA polymerase (6RIC and 6RUI (*36*)) structure being equivalent to the subunits identified by our MS-based proteomics of purified Mimivirus genomic fibers. D] Surface representation of RNA polymerase subunits from vaccinia virus (in cyan), manually docked into the 5-start helical Cl1a map. Each strand of the protein shell is represented in specific color and the dsDNA strands map is shown in pale gold.

## Discussion

Several DNA compaction solutions have been described. For instance, the DNA of filamentous viruses infecting archaea is wrapped by proteins to form a ribbon which in turn folds into a helical rod forming a cavity in its lumen (*37, 38*). In contrast, the chromatin of cellular eukaryotes consists of DNA wrapped around histone complexes (*39*). It was recently shown that the virally encoded histone doublets of the *Marseilleviridae* can form nucleosomes (*40, 41*) and such organization would be consistent with previous evolutionary hypotheses linking giant DNA viruses with the emergence of the eukaryotic nucleus (*42–46*). Herpesviruses (*47, 48*), bacteriophages (*49, 50*) and APBV1 archaeal virus (*35*) package their dsDNA genome as naked helices or coils. APBV1 capsid structure appears strikingly similar to the Mimivirus genomic fiber but Mimivirus, further bundles its flexible genomic fiber as a ball of yarn into the nucleoid, itself encased in the large icosahedral capsids. The structure of the Mimivirus genomic fiber described herein supports a complex assembly process where the DNA must be folded into 5 or 6 strands prior to or concomitant with packaging, a step that may involve the repeat containing regulator of chromosome condensation (qu_366) identified in the proteomic analysis of the genomic fiber. The proteinaceous shell, *via* contacting residues between the dsDNA and the GMC-oxidoreductases, would guide the folding of the dsDNA strands into the structure prior loading into the nucleoid. The lumen of the fiber being large enough to accommodate the Mimivirus RNA polymerase, we hypothesize that it could be sitting on the highly conserved promoter sequence of early genes (*51*). This central position would support the involvement of the RNA polymerase in genome packaging into the nucleoid and could determine the channel width *via* its anchoring on the genome (Fig. 4D). According to this scenario, the available space (although tight) for the RNA polymerase inside the genomic fiber lumen suggests it could be sterically locked inside the compact form of the genomic fiber and could start moving and transcribing upon helix relaxation, initiating the replicative cycle and the establishment of the cytoplasmic viral factory. The genome and the transcription machinery would thus be compacted together into a proteinaceous shield, ready for transcription upon relaxation (Video S3). This organization would represent a remarkable evolutionary strategy for packaging and protecting the viral genome, in a state ready for immediate transcription upon unwinding in the host cytoplasm. This is conceptually reminiscent of icosahedral and filamentous dsRNA viruses which pack and protect their genomes together with the replicative RNA polymerase into an inner core (*52–54*). As a result, replication and transcription take place within the protein shield and viral genomes remain protected during their entire infectious cycle. In the case of dsDNA viruses however, the double helix must additionally open up to allow transcription to proceed, possibly involving the helicase identified in our proteomic study (Table S2). Finally, in addition to their structural roles, the FAD containing GMC-oxidoreductases making the proteinaceous shield, together with other oxidative stress proteins (Table S2), could alleviate the oxidative stress to which the virions are exposed while entering the cell by phagocytosis. The complexity of the flexible genomic fiber, with various relaxation states requires additional work aiming to stabilize this macromolecular complex in additional intermediate states in order to determine the contributions of the various proteins identified and ultimately to progress towards structures of the process inside host cells in the course of Mimivirus infection.

Mimivirus virion thus appears as a Russian doll, with its icosahedral capsids covered with heavily glycosylated fibrils, two internal membranes, one lining the capsid shell, the other encasing the nucleoid, in which the genomic fiber is finally folded. To our knowledge, the structure of the genomic fiber used by Mimivirus to package and protect its genome in the nucleoid represents the first description of the genome organization of a giant virus. Since the genomic fiber appears to be expelled from the nucleoid as a flexible and subsequently straight structure starting decompaction upon release, we suspect that an active, energy-dependent, process is required to bundle it into the nucleoid during virion assembly. Such an efficient structure is most likely shared by other members of the Mimiviridae family infecting *Acanthamoeba* and could be used by other dsDNA viruses relying on exclusively cytoplasmic replication like poxviruses to immediately express early genes upon entry into the infected cell (*55, 56*). Finally, the parsimonious use of moonlighting GMC-oxidoreductases playing a central role in two functionally unrelated substructures of the Mimivirus particle - i) as a component of the heavily glycosylated peripheral fibril layer and ii) as a proteinaceous shield to package the dsDNA into the genomic fibers questions the evolutionary incentive leading to such an organization for a virus encoding close to a thousand proteins.

## Materials and Methods

### Nucleoid extraction

Mimivirus reunion virions defibrillated, as described previously (*11, 19*), were centrifuged at 10,000 x g for 10 min, resuspended in 40 mM TES pH 2 and incubated for 1h at 30°C to extract the nucleoid from the opened capsids.

### Extraction and purification of the Mimivirus genomic fiber

The genomic fiber was extracted from 12 mL of purified Mimivirus reunion virions at 1.5 x 10^10^ particles/mL, split into 12×1 mL samples processed in parallel. Trypsin (Sigma T8003) in 40 mM Tris-HCl pH 7.5 buffer was added at a final concentration of 50 µg/mL and the virus-enzyme mix was incubated for 2h at 30°C in a heating dry block (Grant Bio PCH-1). DTT was then added at a final concentration of 10 mM and incubated at 30°C for 16h. Finally, 0.001% Triton X-100 was added to the mix and incubated for 4h at 30°C. Each tube was vortexed for 20 s with 1.5 mm diameter stainless steel beads (CIMAP) to separate the fibers from the viral particles and centrifuged at 5,000 x g for 15 min to pellet the opened capsids. The supernatant was recovered, and the fibers were concentrated by centrifugation at 15,000 x g for 4h at 4°C. Most of the supernatant was discarded leaving 12x∼200 µL of concentrated fibers that were pooled and layered on top of ultracentrifuge tubes of 4 mL (polypropylene centrifuge tubes, Beckman Coulter) containing a discontinuous sucrose gradient (40%, 50%, 60%, 70% w/v in 40 mM Tris-HCl pH 7.5 buffer). The gradients were centrifuged at 200,000 x g for 16h at 4 °C. Since no visible band was observed, successive 0.5 mL fractions were recovered from the bottom of the tube, the first one supposedly corresponding to 70% sucrose. Each fraction was dialyzed using 20 kDa Slide-A-Lyzers (ThermoFisher) against 40 mM Tris-HCl pH 7.5 to remove the sucrose. These fractions were further concentrated by centrifugation at 15,000 x g, at 4°C for 4h and most of the supernatant was removed, leaving ∼100 µL of sample at the bottom of each tube. At each step of the extraction procedure the sample was imaged by negative staining transmission electron microscopy (TEM) to assess the integrity of the genomic fiber (Fig. S1). Each fraction of the gradient was finally controlled by negative staining TEM. For proteomic analysis, an additional step of concentration was performed by speedvac (Savant SPD131DDA, Thermo Scientific).

### Negative stain TEM

300 mesh ultra-thin carbon-coated copper grids (Electron Microscopy Sciences, EMS) were prepared for negative staining by adsorbing 4-7 µL of the sample for 3 min., followed by two washes with water before staining for 2 min in 2% uranyl acetate. The grids were imaged either on a FEI Tecnai G2 microscope operated at 200 keV and equipped with an Olympus Veleta 2k camera (IBDM microscopy platform, Marseille, France); a FEI Tecnai G2 microscope operated at 200 keV and equipped with a Gatan OneView camera (IMM, microscopy platform, France) or a FEI Talos L120c operated at 120 keV and equipped with a Ceta 16M camera (CSSB multi-user cryo-EM facility, Germany, Fig. 1, Fig. S1).

### Agarose gel electrophoresis and DNA dosage to assess the presence of DNA into the fiber

Genomic DNA was extracted from 10^10^ virus particles using the PureLink TM Genomic DNA mini kit (Invitrogen) according to the manufacturer’s protocol. Purified genomic fiber was obtained following the method described above. The purified fiber was treated by adding proteinase K (PK) (Takara ST 0341) to 20 µL of sample (200 ng as estimated by dsDNA Qubit fluorometric quantification) at a final concentration of 1 mg/mL and incubating the reaction mix at 55°C for 30 min. DNase treatment was done by adding DNase (Sigma 10104159001) and MgCl_2_ to a final concentration of 0.18 mg/mL and 5 mM, respectively, in 20 µL of sample and incubated at 37°C for 30 min prior to PK treatment. For RNase treatment, RNase (Sigma SLBW2866) was added to 20 µL (200 ng) of sample solution to a final concentration of 1 mg/mL and incubated at 37°C for 30 min prior to PK treatment. All the samples were then loaded on a 1 % agarose gel and stained with ethidium bromide after migration. The bands above the 20 kbp marker correspond to the stacked dsDNA fragments of various lengths compatible with the negative staining images of the broken fibers where long DNA fragment are still attached to the helical structure. The Mimivirus purified genomic DNA used as a control migrates at the same position (Fig. S2).

### Cryo-EM bubblegram analysis

Samples were prepared as described for single-particle analysis. Dose series were acquired on a Titan Krios (Thermo Scientific) microscope operated at 300 keV and equipped with a K3 direct electron detector and a GIF BioQuantum (Gatan) energy filter. Micrographs were recorded using SerialEM (*57*) at a nominal magnification of 81,000x, a pixel size of 1.09 Å and a rate of 15 e^-^/pixel/s (Fig. S3 & Table S3). Dose series were acquired by successive exposures of 6 s, resulting in an irradiation of 75 e^-^/Å² per exposure. Micrographs were acquired with 0.1 s frames and aligned in SerialEM (*57*). In a typical bubblegram experiment, 12 to 15 successive exposures were acquired in an area of interest with cumulative irradiations of 900 to 1,125 e^-^/Å² total (Fig. S3).

### Single-particle analysis by cryo-EM

#### Sample preparation

For single-particle analysis, 3 µL of the purified sample were applied to glow-discharged Quantifoil R 2/1 Cu grids, blotted for 2 s using a Vitrobot Mk IV (Thermo Scientific) and applying the following parameters: 4°C, 100% humidity, blotting force 0, and plunge frozen in liquid ethane/propane cooled to liquid nitrogen temperature.

#### Data acquisition

Grids were imaged using a Titan Krios (Thermo Scientific) microscope operated at 300 keV and equipped with a K2 direct electron detector and a GIF BioQuantum energy filter (Gatan). 7,656 movie frames were collected using the EPU software (Thermo Scientific) at a nominal magnification of 130,000x with a pixel size of 1.09 Å and a defocus range of −1 to −3 μm. Micrographs were acquired using EPU (Thermo Scientific) with 8 s exposure time, fractionated into 40 frames and 7.5 e^-^/pixel/ s (total fluence of 50.5 e^-^/Å²) (Table S3).

#### 2D classification and clustering of 2D classes

All movie frames were aligned using MotionCor2 (*58*) and used for contrast transfer function (CTF) estimation with CTFFIND-4.1 (*59*). Helical segments of the purified genomic fibers, manually picked with Relion 3.0 (*28, 29*)(*28, 29*), were initially extracted with different box sizes, 400 pixels for 3D reconstructions, 500 pixels for initial 2D classifications and clustering and 700 pixels to estimate the initial values of the helical parameters. Particles were subjected to reference-free 2D classification in Relion 3.1.0 (*28, 29*), where multiple conformations of the fiber were identified (Fig. S4, S5).

We then performed additional cluster analysis of the 194 initial 2D classes provided by Relion (Fig. S4) to aim for more homogeneous clusters, eventually corresponding to different states (Fig. S5). A custom 2-step clustering script was written in python with the use of Numpy(*60*) and Scikit-learn (*61*) libraries. First, a few main clusters were identified by applying a DBSCAN (*62*) clustering algorithm on the previously estimated fiber external width values (W1). The widths values, estimated by adjusting a parameterized cross-section model on each 2D stack, range from roughly 280 Å to 340 Å. The cross-section model fitting process is based on adjusting a section profile (S) described in Equation 1 on the longitudinally integrated 2D-class profile (Fig. S5A). The cross-section model is composed of a positive cosine component, (parameterized by the center position μ, its width σ_1_ and amplitude a_1_) associated with the fiber external shell, a negative cosine component (parameterized by the center position μ, its width σ_2_ and amplitude a_2_) associated with the central hollow lumen (W2), and a constant p_0_, accounting for the background level, as

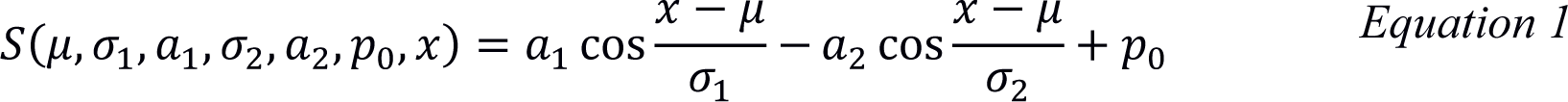

Then, as a second step, each main cluster was subdivided into several sub-clusters by applying a KMEANS(*63*) clustering algorithm on a pairwise similarity matrix. This similarity metric was based on a 2D image cross-correlation scheme, invariant to image shifts, and mirroring (*64*). The number of sub-clusters was manually chosen by visual inspection. For the most populated 2D classes corresponding to the most compact conformations, the number of sub-clusters was small: n=2 sub-clusters, Cl1a and Cl1b, and n=1 sub-cluster for the intermediate class, Cl2. However, the number of sub-clusters was higher (n=5) for the relaxed conformation (Cl3 from green to purple in Fig. S5), highlighting the overall heterogeneity of our dataset with compact, intermediate, relaxed states and even loss of one protein strands (Cl3b) and unwound ribbons.

### Identification of candidate helical parameters

Fourier transform analysis methods have been used to identify helical parameters candidates (*65–67*) for the Cl1a, Cl2 and Cl3a clusters. For each cluster, we first estimated the repeat distance by applying the method consisting in a longitudinal autocorrelation with a windowed segment of the real space 2D class of fixed size (100 Å) (*66*). Then, a precise identification of the power spectrum maxima could be achieved on a high signal-to-noise ratio power spectrum, obtained by averaging all the constituting segments in the Fourier domain, which helps lowering the noise, and fill in the CTF zeros regions. The best candidates were validated with Helixplorer (http://rico.ibs.fr/helixplorer/).

For the Cl1a cluster, the parameters of a 1-start helix have been identified with a rise of 7.937 Å and a twist of 221.05°. For the Cl2 cluster, the candidate parameters are a rise of 20.383 Å and a twist of 49.49° and C3 cyclic symmetry. For the Cl3a cluster, the candidate parameters correspond to a rise of 31.159 Å and a twist of 24° and D5 symmetry.

### Cryo-EM data processing and 3D reconstruction

–Cl1a-Cl3a After helical parameters determination, two last 2D classifications were performed on segments extracted with a box size of 400 pixels (decimated to 200 pixels) using the proper rises for the most compact Cl1a (7.93 Å, 113,026 segments, overlap ∼98.2%) and the relaxed Cl3a (31.16 Å, 16,831 segments, overlap ∼92.9%). Values of the helical parameters (rise and twist) were then used for Relion 3D classification (*28, 29*), with a +/- 10% freedom search range, using a featureless cylinder as initial reference (diameter of 300 Å for the compact particles Cl1a and 340 Å for the relaxed particles Cl3a). The superimposable 3D classes (same helical parameters, same helix orientation) were then selected, reducing the data set to 95,722 segments for the compact fiber (Cl1a) and to 15,289 segments for the relaxed fiber (Cl3a). After re-extraction of the selected segments without scaling, further 3D refinement was performed with a 3D classification output low pass filtered to 15 Å as reference. With this, the maps were resolved enough (Cl1a: 4.4 Å, Cl3a: 4.8; FSC threshold 0.5) to identify secondary structure elements (with visible cylinders corresponding to the helices) (Fig. S6-7).
–Cl2 The twelve 2D classes corresponding to segments of the Cl2 conformation, were extracted with a box size of 400 pixels (decimated to 200 pixels, rise 20.4 Å, 5,775 segments, overlap ∼94.9%). They were used in Relion for 3D refinement with the helical parameters values identified previously, with a +/-10% freedom search range, using a 330 Å large featureless cylinder as initial reference and resulted in a 7.1 Å map (FSC threshold 0.5) and C3 cyclic symmetry (Fig. S8).

### Focused refinement of the asymmetric unit

The Cl1a map, prior to solvent flattening, was used to obtain the mask corresponding to a single asymmetric unit of the structure (just one dimer). It was then used to subtract everything else from the corresponding experimental images, keeping only information from that asymmetric unit for a following round of focused refinement. The subtracted dataset was then used as input for Relion in order to reconstruct a reference for 3D refinement of the asymmetric unit with a last solvent flattening step. Further CTF refinement steps were performed before last 3D refinement with solvent flattening and post-processing (B-factor applied −45). This led to the best resolved 3.3 Å map (FSC threshold 0.5, masked, Fig. S6) that was used to refine the GMC-oxidoreductase dimer and its ligands (Table S1, Fig. 3). While the mask was only based on the protein asymmetric unit the DNA contribution is still visible in the final map.

### Automatic picking of the three different conformations (Cl1a, Cl2, Cl3a)

Projections from the different helical maps were used as new input for automatic picking of each Cl1a, Cl2 and Cl3a clusters to get more homogeneous datasets for each conformation (*68*). For each dataset a final round of extraction (box size Cl1a: 380 pixels, 121,429 segments; Cl2: 400 pixels, 8479 segments; Cl3a: 400 pixels, 11,958 segments) and 3D refinement with solvent flattening (Central z length 30%) was performed using the appropriate helical parameters and additional symmetries (none for Cl1a, C3 for Cl2 and D5 for Cl3a). This led to the improved maps presented in Fig. 2-4 and Fig. S6-S8 (Cl1a: 3.7 Å; Cl2: 4.0 Å; Cl3a: 3.7 Å; FSC threshold 0.5, masked). Post-processing was performed with B-factor −80 (Table S1, Fig. S6-S8).

### DNA focus refinement

The Cl1a and Cl2 maps without solvent flattening were used to obtain the mask corresponding to each DNA strand independently. Each strand was then used for subtraction keeping only information from the corresponding DNA on the corresponding experimental images. The subtracted datasets of each DNA strand were then merged. The resulting dataset was used to perform 2D classifications in order to assess the quality of the subtracted dataset (Fig. S9). It was then used to perform a 3D refinement followed by a 3D classification and final refinement of the best 3D classes (Fig. 3E, Fig. S10).

For the Cl3a compaction states, a cylinder with a diameter corresponding to the lumen of the relaxed structure was used as mask for subtraction, in order to try to enhance the information corresponding to the sole DNA. However, the DNA signal remained too weak.

### Structures refinement

The resolution of the EM map enabled to fit the R135 dimeric structure (*30*) (PDB 4Z24) into the maps using UCSF Chimera 1.13.1 (*69*). The qu_143 and qu_946 models were obtained using SWISS-MODEL (*70*) (closest PDB homologue: 4Z24). It is only at that stage that the best fitted qu_946 model was manually inspected and additional N-terminal residues built using the extra density available in the cryo-EM map of the 5-start compact reconstruction (Fig. 3). The entire protein shell built using the corresponding helical parameters were finally fitted into the Cl1a and Cl3a maps and were further refined against the map using the real-space refinement program in PHENIX 1.18.2 (*33*) (Fig. S6-7). The qu_143 model was selected to build the entire shell using the Cl2 helical parameters and symmetry and was further refined using the real-space refinement program in PHENIX 1.18.2 (*33*) (Fig. S8). Validations were also performed into PHENIX 1.18.2 (*33*) using the comprehensive validation program (Table S1). The qu_946 model was manually corrected and ultimately refined and validated into PHENIX 1.18.2 (*33*) using the highest resolution focused refined Cl1a map. In that map the 2 first amino acid are disordered including the cysteine.

### Cryo-electron tomography

#### Sample preparation

For cryo-ET of the Mimivirus genomic fiber, samples were prepared as described above for single-particle analysis except that 5 nm colloidal gold fiducial markers (UMC, Utrecht) were added to the sample right before plunge freezing at a ratio of 1:2 (sample: fiducial markers).

#### Data acquisition

Tilt series were acquired using SerialEM (*57*) on a Titan Krios (Thermo Scientific) microscope operated at 300 keV and equipped with a K3 direct electron detector and a GIF BioQuantum energy filter (Gatan). We used the dose-symmetric tilt-scheme (*71*) starting at 0° with a 3° increment to +/-60° at a nominal magnification of 64,000x, a pixel size of 1.4 Å and a total fluence of 150 e-/Å² over the 41 tilts (*i.e.* ∼3.7 e-/Å²/tilt for an exposure time of 0.8 s fractionated into 0.2 s frames (Table S3).

#### Data processing

Tilt series were aligned and reconstructed using the IMOD (*72*). For visualization purposes, we applied a binning of 8 and SIRT-like filtering from IMOD (*72*) as well as a bandpass filter bsoft (*73*). The tomograms have been deposited on EMPIAR, accession number 1131 and videos were prepared with Fiji (Fig. 1E-F, Fig. S11 & Tomo. S1-S4).

### Mass spectrometry-based proteomic analysis of Mimivirus virion and genomic fiber

Proteins extracted from total virions and purified fiber were solubilized with Laemmli buffer (4 volumes of sample with 1 volume of Laemmli 5X - 125 mM Tris-HCl pH 6.8, 10% SDS, 20% glycerol, 25% β-mercaptoethanol and traces of bromophenol blue) and heated for 10 min at 95 °C. Three independent infections using 3 different batches of virions were performed and the genomic fiber was extracted from the resulting viral particles to analyse three biological replicates. Extracted proteins were stacked in the top of a SDS-PAGE gel (4-12% NuPAGE, Life Technologies), stained with Coomassie blue R-250 (Bio-Rad) before in-gel digestion using modified trypsin (Promega, sequencing grade) as previously described (*74*). Resulting peptides were analyzed by online nanoliquid chromatography coupled to tandem MS (UltiMate 3000 RSLCnano and Q-Exactive Plus, Thermo Scientific). Peptides were sampled on a 300 µm x 5 mm PepMap C18 precolumn and separated on a 75 µm x 250 mm C18 column (Reprosil-Pur 120 C18-AQ, 1.9 μm, Dr. Maisch) using a 60-min gradient for fiber preparations and a 140-min gradient for virion. MS and MS/MS data were acquired using Xcalibur (Thermo Scientific). Peptides and proteins were identified using Mascot (version 2.7.0) through concomitant searches against Mimivirus reunion, classical contaminant databases (homemade) and the corresponding reversed databases. The Proline software (*75*) was used to filter the results: conservation of rank 1 peptides, peptide score ≥ 25, peptide length ≥ 6, peptide-spectrum-match identification false discovery rate < 1% as calculated on scores by employing the reverse database strategy, and minimum of 1 specific peptide per identified protein group. Proline was then used to perform a compilation and MS1-based quantification of the identified protein groups. Intensity-based absolute quantification (iBAQ) (*76*) values were calculated from MS intensities of identified peptides. The viral proteins detected in a minimum of two replicates are reported with their molecular weight, number of identified peptides, sequence coverage and iBAQ values in each replicate. The number of copies per fiber for each protein was calculated according to their iBAQ value based on the GMC-oxidoreductases iBAQ value (see below, Table S2). Mapping of the identified peptides for the GMC-oxidoreductases are presented in Fig. S12.

### Protein/DNA ratio validation

To compare the theoretical composition of the genomic fiber accommodating a complete genome with the experimental concentrations in protein and DNA of the sample, we performed DNA and protein quantification (Qubit fluorometric quantification, Thermo Fischer Scientific) on two independent samples of the purified genomic fiber. This returned a concentration of 1.5 ng/µL for the dsDNA and 24 ng/µL for the proteins in one sample and 10 ng/µL for the dsDNA and 150 ng/µL for the protein in the second sample (deposited on agarose gel, Fig. S2). Considering that all proteins correspond to the GMC-oxidoreductases subunits (∼71 kDa), in the sample there is a molecular ratio of 7.2 dsDNA base pairs per GMC-oxidoreductases subunit in the first sample and 7.68 in the second. Based on our model, a 5-start genomic fiber containing the complete Mimivirus genome (1,196,989 bp) should be composed of ∼95,000 GMC-oxidoreductases subunits (∼93,000 for a 6-start). This gives a molecular ratio of 12.5 dsDNA base pairs per GMC-oxidoreductase subunit, which would be more than experimentally measured. However, in the cryo-EM dataset, there are at most 50% of genomic fibers containing the DNA genome (mostly Cl1), while some DNA strands can be observed attached to the relaxed genomic fiber Cl3 but the vast majority of released DNA was lost during purification of the genomic fiber. Applying an estimated loss of 50 % to the total DNA compared to the measured values, we obtain a ratio, which is in the same order of magnitude of the ones measured in the purified genomic fiber samples.

### Model visualization

Molecular graphics and analyses were performed with UCSF Chimera 1.13.1 (*69*) and UCSF ChimeraX 1.1 (*77*), developed by the Resource for Biocomputing, Visualization, and Informatics at the University of California, San Francisco, with support from National Institutes of Health R01-GM129325 and the Office of Cyber Infrastructure and Computational Biology, National Institute of Allergy and Infectious Diseases.

## Acknowledgements

The cryo-EM work was performed at the multi-user Cryo-EM Facility at CSSB. We thank Jean-Michel Claverie for his comments on the manuscript and discussions all along the project. We thank Irina Gutsche, Ambroise Desfosses, Eric Durand and Juan Reguera for their helpful support on structural work. We thank Carolin Seuring for support and technical help. Processing was performed on the DESY Maxwell cluster, at IBS and IGS. We thank Wolfgang Lugmayr for assistance and Sebastien Santini and Guy Schoehn for support. The preliminary electron microscopy experiments were performed on the PiCSL-FBI core facility (Nicolas Brouilly, Fabrice Richard and Aïcha Aouane, IBDM, AMU-Marseille), member of the France-BioImaging national research infrastructure and on the IMM imaging platform (Artemis Kosta).

## Authors contributions

Conceptualization and Project administration: CA Methodology: AV, AS, LFE, EQ, JMA, AL, LB, YC and CA Formal Analysis: AV, AS, LFE, EQ, JMA, AL, LB, YC and CA Software: AS, AV, LFE, CA Investigation: AV, AS, LFE, EQ, JMA, AL, LB, AC, FH, YC and CA Funding acquisition: CA, KG, EQ, YC Resources: KG granted access to acquisition and analytic tools Writing – original draft: CA Writing – review & editing: AV, AS, EQ, KG and CA

## Data and materials availability

Mimivirus reunion genome has been deposited under the following accession number: BankIt2382307 Seq1 MW004169. 3D reconstruction maps and the corresponding PDB have been deposited to EMDB (Deposition number Cl1a: 7YX4, EMD-14354; Cl1a focused refined: 7PTV, EMD-13641; Cl3a: 7YX5, EMD-14355; Cl2: 7YX3, EMD-14353). The mass spectrometry proteomics data have been deposited to the ProteomeXchange Consortium via the PRIDE partner repository with the dataset identifier PXD021585 and 10.6019/PXD021585. The tomograms have been deposited in EMPIAR under the accession number 1131.

## Supplementary Materials

The Videos, tomograms, EM maps and pdb files can be found under the following link: https://s.42l.fr/mfiberdata

---

**Fig. S1:**
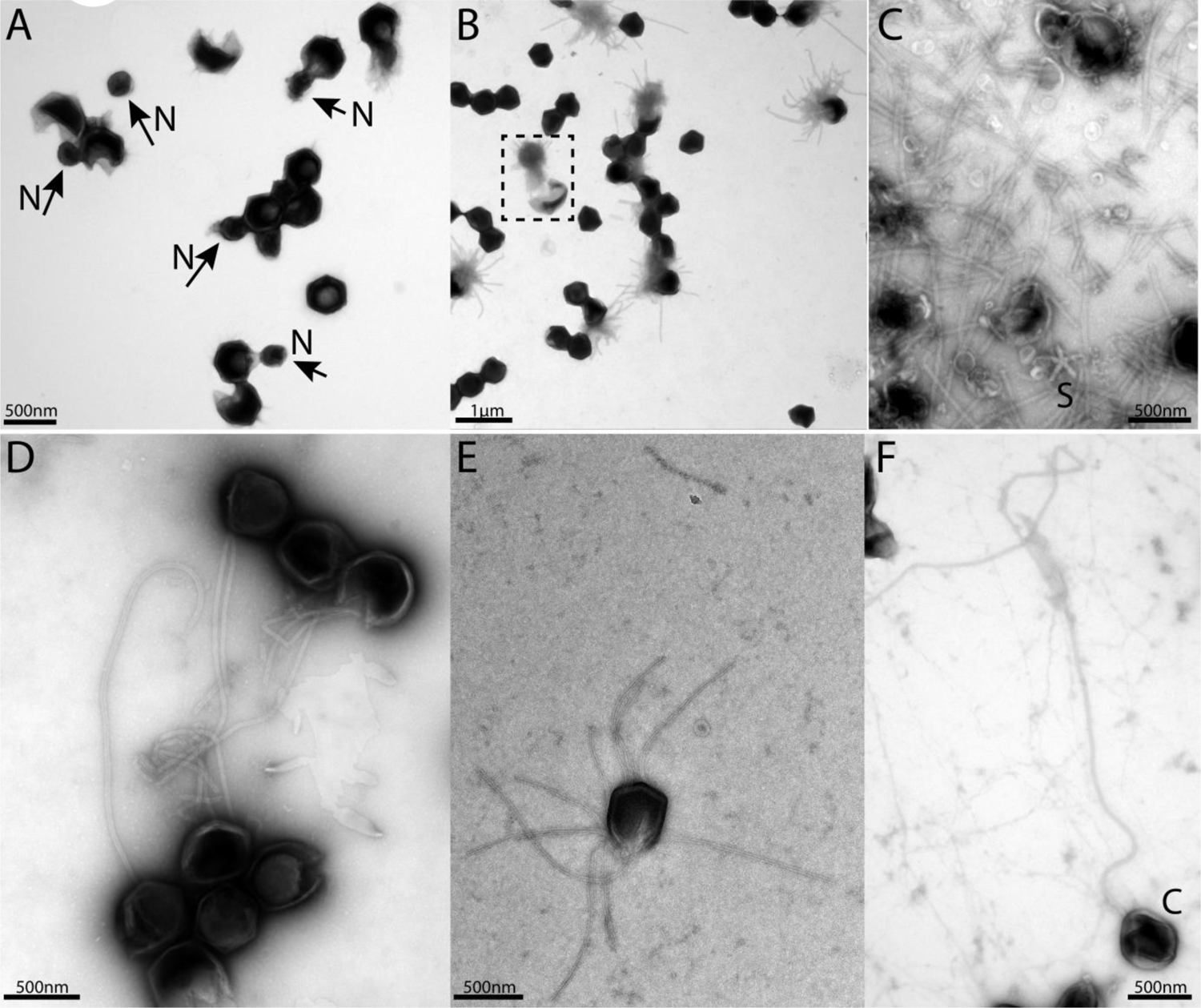
Negative staining micrographs of opened Mimivirus reunion virions before purification of the genomic fiber. (**A**) Spherical nucleoids (N) released from the capsids upon specific treatment (see methods). (**B**) Dissolved nucleoids expelled from the opened capsids after regular treatment (the region marked with dashed lines is enlarged in Fig 1C), all of the opened capsids (>25%) are releasing the condensed genomic fiber. (**C**) Multiple broken genomic fibers can be seen next to debris prior to purification and an isolated Stargate (S). (**D-E**) Multiple segments of the flexible genomic fiber are released upon proteolytic treatment of the capsids. (**F**) Example of a long, flexible, genomic fiber released from an open capsid (C).

**Fig. S2:**
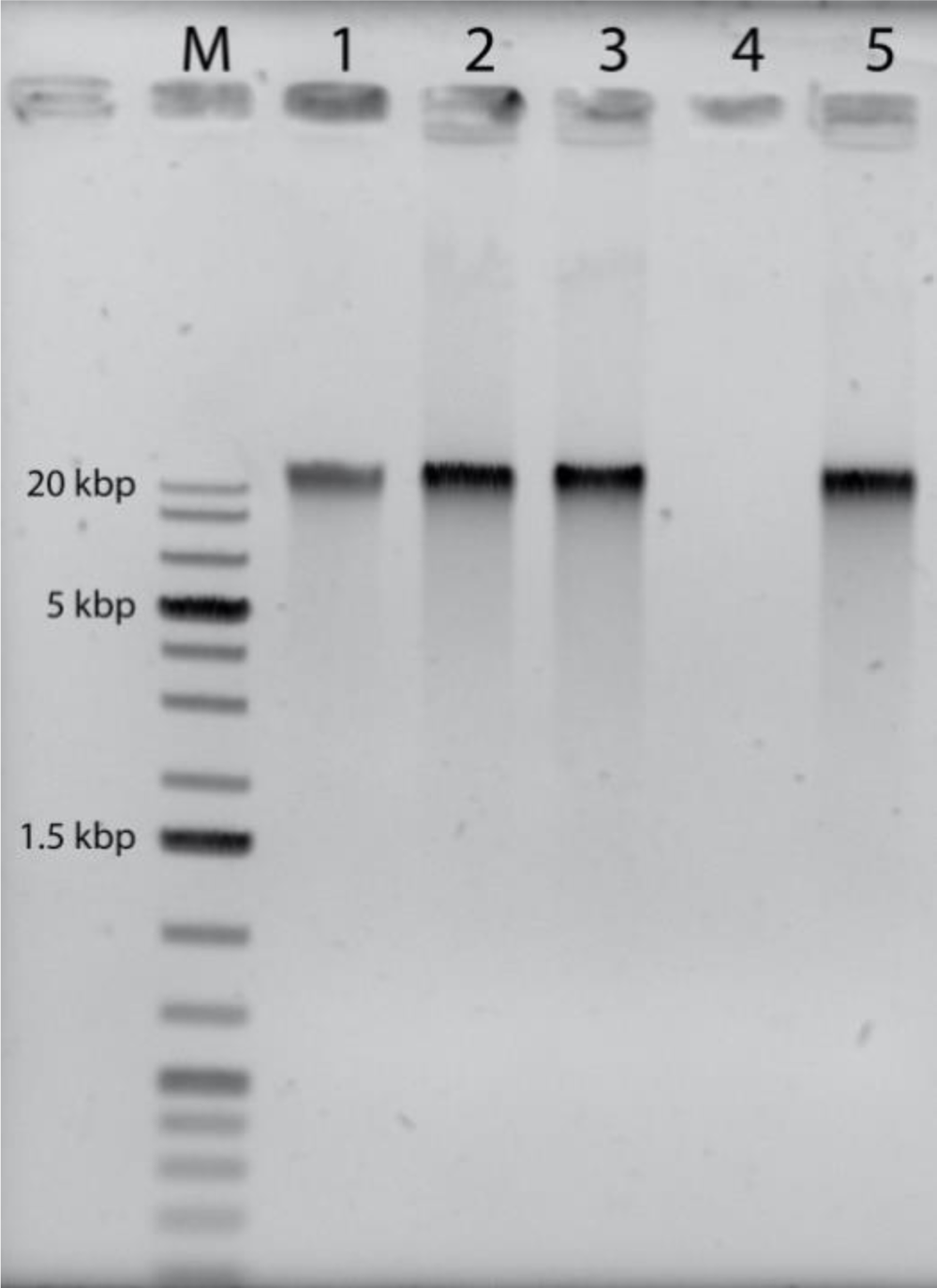
Agarose gel electrophoresis of the purified genomic fiber compared to the viral genomic DNA: M: Molecular weight markers (1 kbp DNA Ladder Plus, Euromedex). Lane 1, 100 ng of Mimivirus genomic DNA, Lane 2: 200 ng of untreated genomic fiber, Lane 3: 200 ng of Proteinase K-(PK) treated genomic fiber, Lane 4: 200 ng of DNase- and PK-treated genomic fiber, Lane 5: 200 ng of RNase- and PK-treated genomic fiber.

**Fig. S3:**
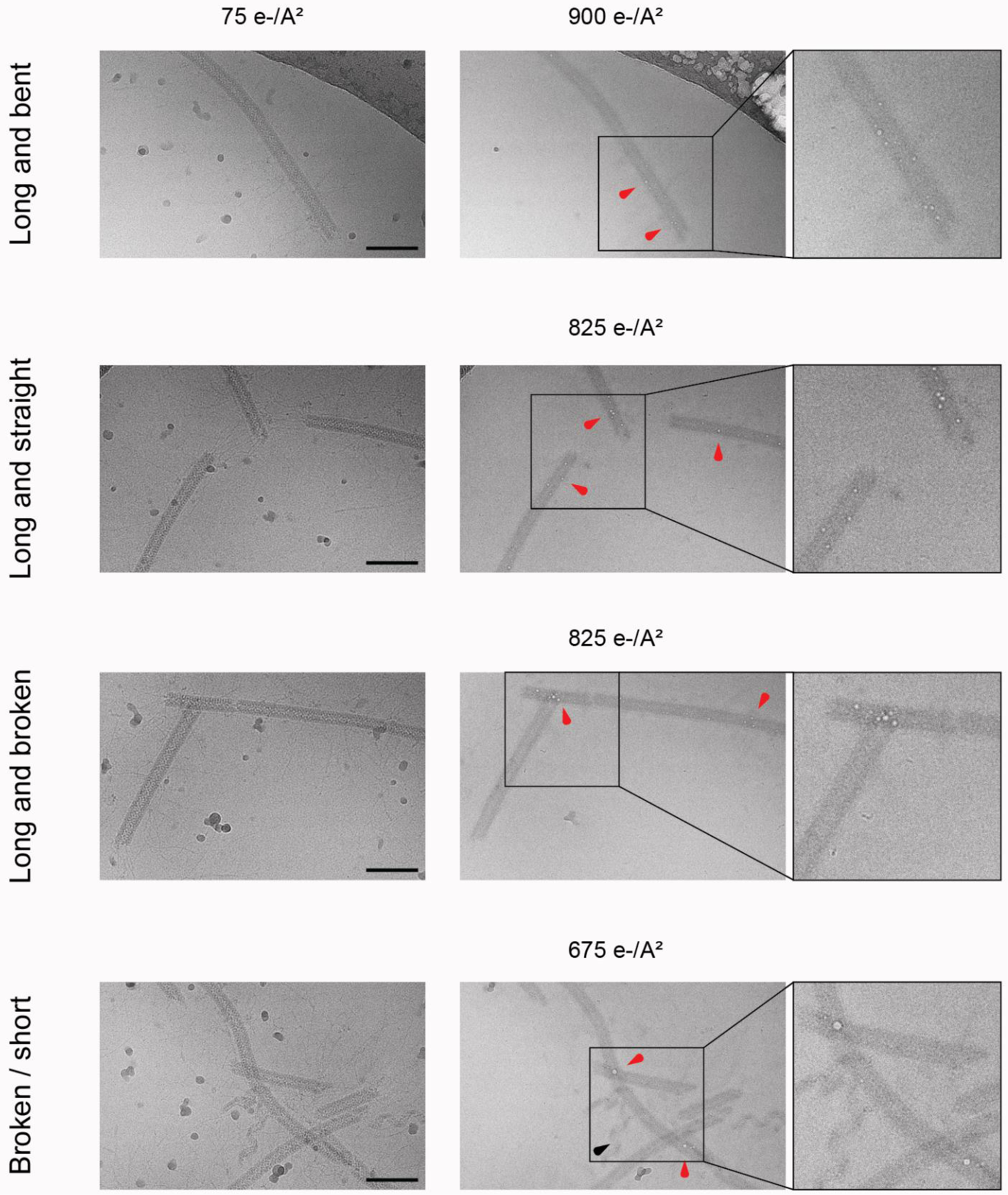
Bubblegrams on the Mimivirus genomic fiber. Field of view for genomic fibers either (from top to bottom): long and bent, long and straight, long and broken, or a mix of short and broken, subjected to a dose series. The total electron flux applied is specified for each field of view. First column: fibers showed after the first exposure of 75 e^-^/Å². Second column: representative micrographs indicating the point at which the first sign of radiation damage as “bubbles” inside the fiber (red arrowheads) were detected. Nucleoproteins with high nucleic acid content are often more susceptible to bubbling than pure proteinaceous structures. In unfolded ribbons (black arrowhead), no bubbles are detected at an accumulated flux of up to 1,125 e-/Å². Scale bars, 100 nm. Third column: enlarged view where the effect of radiation damage as trapped “bubbles” is more clearly visible.

**Fig. S4:**
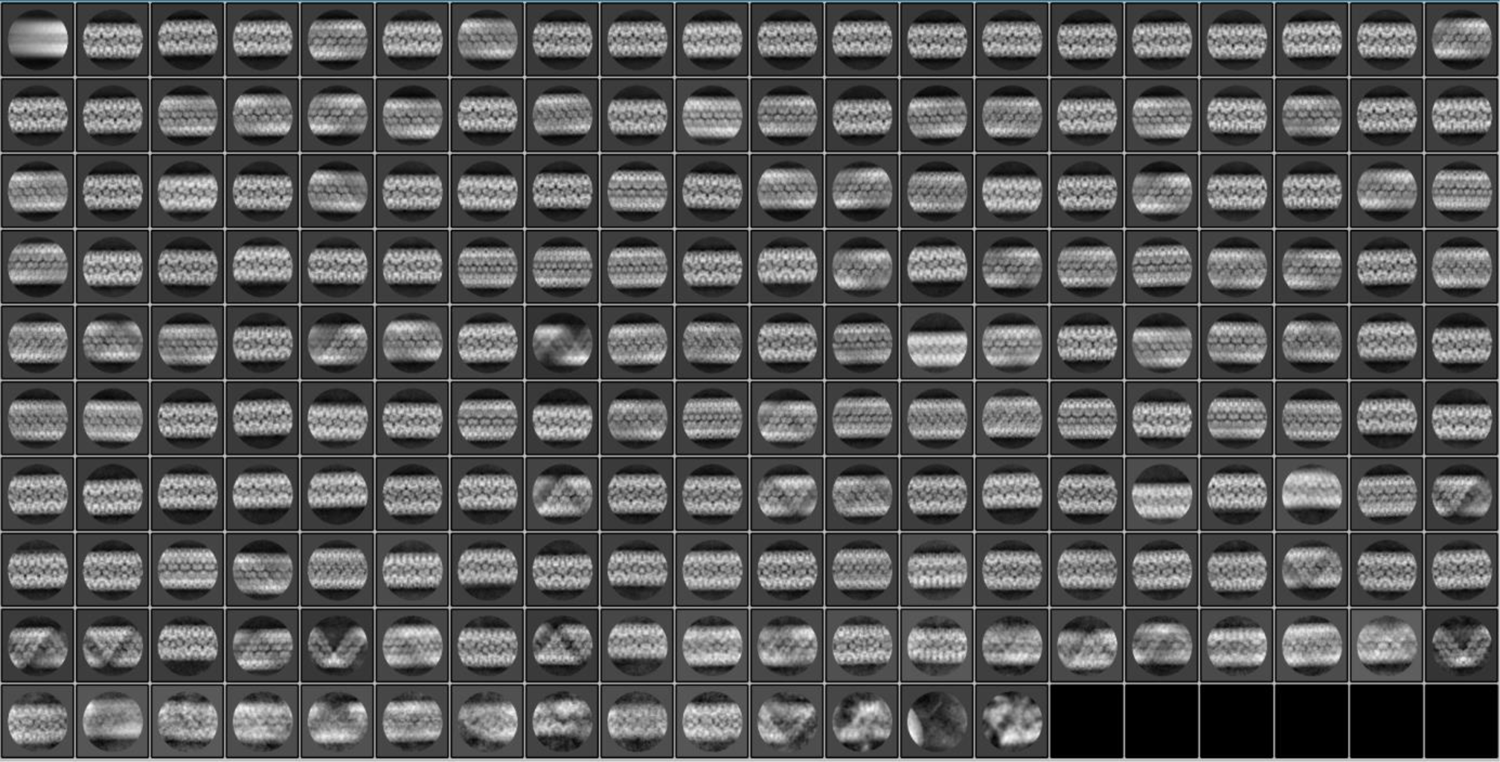
2D classification. 200 2D classes (6 empty) were obtained after reference-free 2D classification of fibers acquired (see methods) for single-particle analysis and extracted with a box size of 500 pixels in Relion 3.1.0 after motion correction, CTF estimation and manual picking. The 2D classes are representative of the different relaxation states of the Mimivirus genomic fiber observed in our highly heterogeneous dataset.

**Fig. S5:**
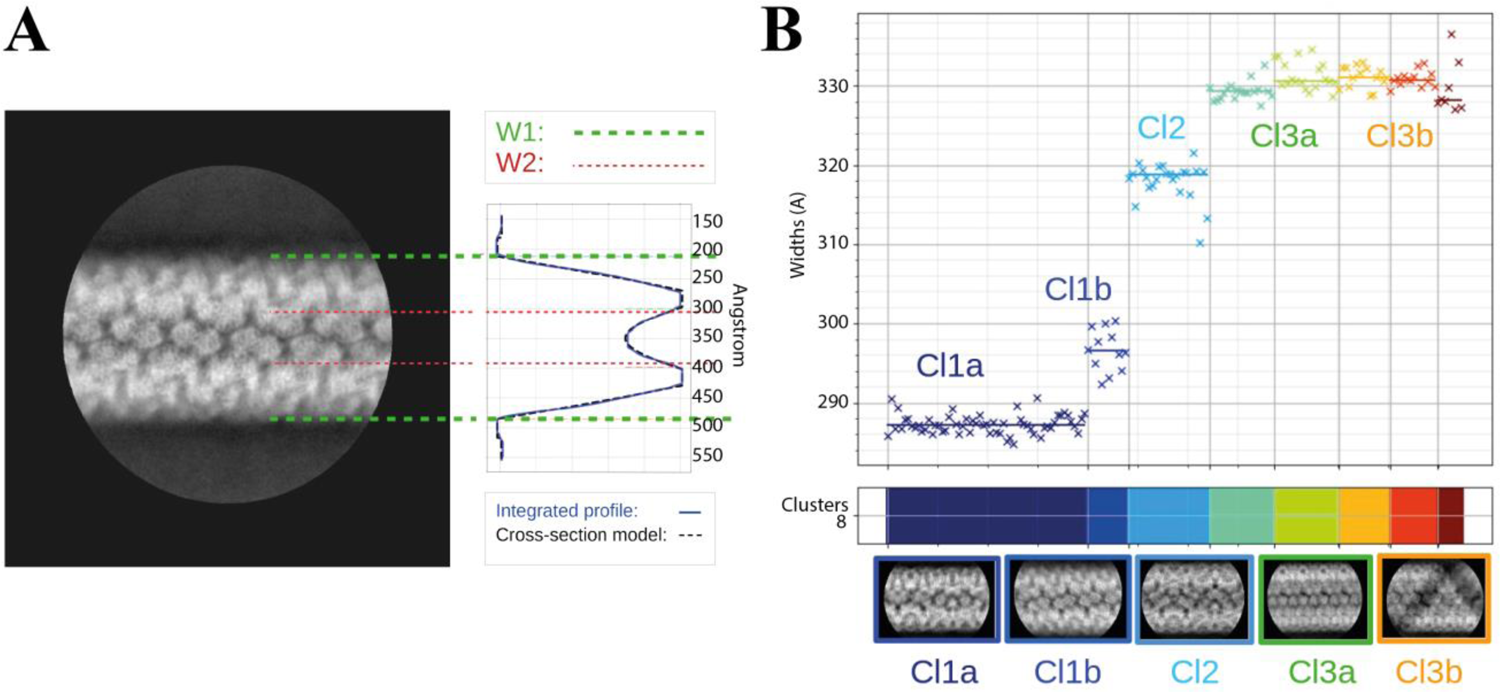
Clustering analysis of the 2D classes. **(A)** Cross-section model adjusted to one 2D class, representative of cluster Cl1b, to estimate the parameters of the widths W1 (external) and W2 (internal). (**B**) Automatic sorting of the 2D classes using the fiber width W1 and pairwise correlations of the 2D classes resulting in 3 main clusters (compact, Cl1 in dark blue; intermediate, Cl2 in cyan and relaxed, Cl3 in green and orange).

**Fig. S6:**
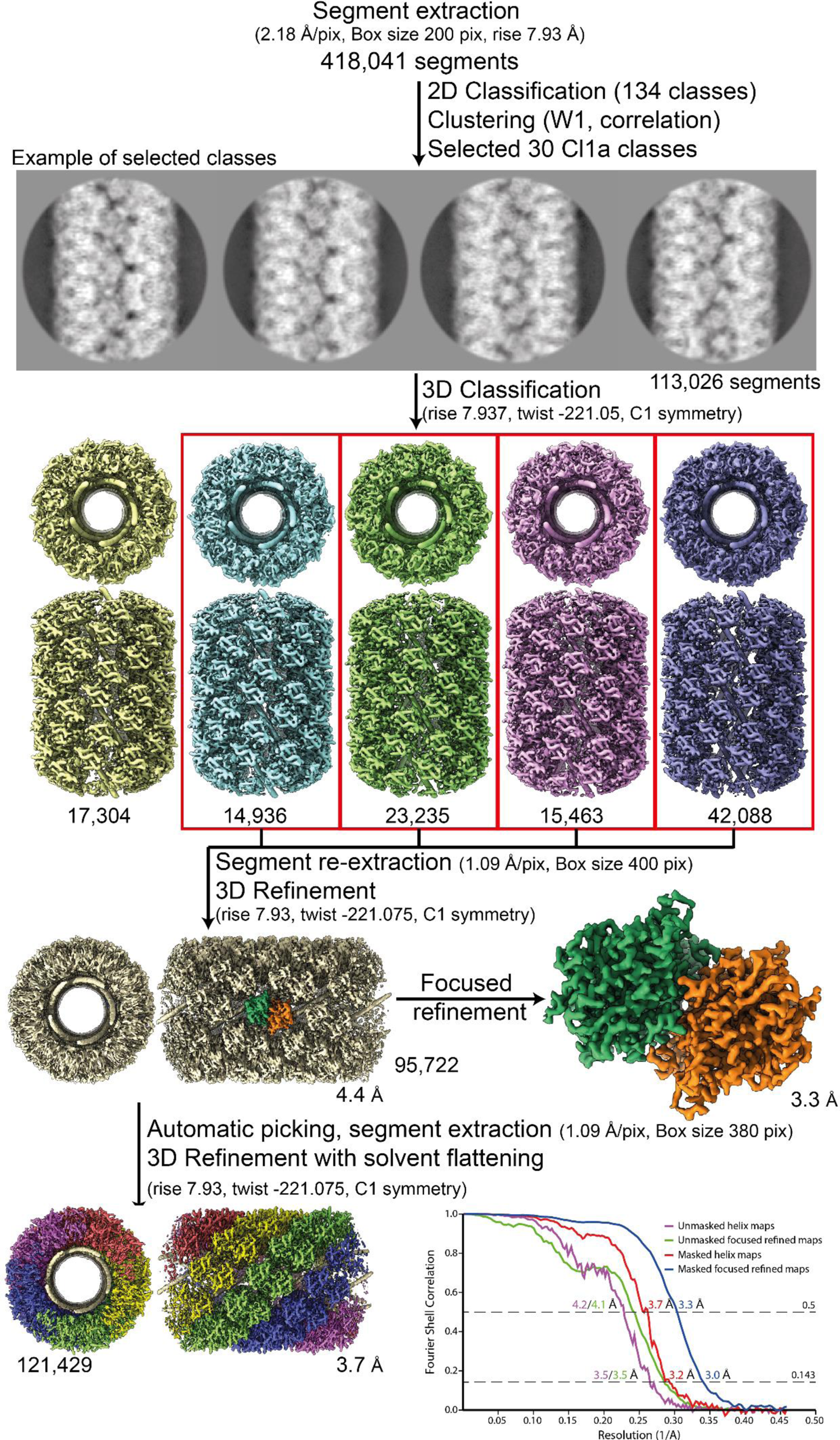
Workflow of the Cl1a compact helix reconstruction process and fitting. Segment extraction was performed with a box size of 400 pixels (pix) binned twice (box size 200 pix, 2.18 Å/pix). The distance between consecutive boxes was equal to the axial rise calculated by indexation on the power spectrum. From 2D classification with 200 classes, 30 classes were selected for Cl1a after clustering (see methods and Fig. S5). First, 3D classification was carried out using the segments from 30 2D classes, helical symmetry parameters from the power spectrum indexation and a 300 Å featureless cylinder as 3D reference. 3D-refinement of the 4 boxed 3D-classes was achieved using one low pass filtered 3D class as reference on the unbinned segments. One step of focused refinement (with solvent flattening) was performed using a low pass filtered 3D reconstruction (before solvent flattening) as reference. 3D-refinement was then performed. The resulting map was then used as model for automatic picking. A last 3D-refinement cycle was performed using as reference a featureless cylinder. The bottom right graph presents the Fourrier shell correlation (FSC) curves for the final 3D reconstructions (helical assembly and asymmetric unit).

**Fig. S7:**
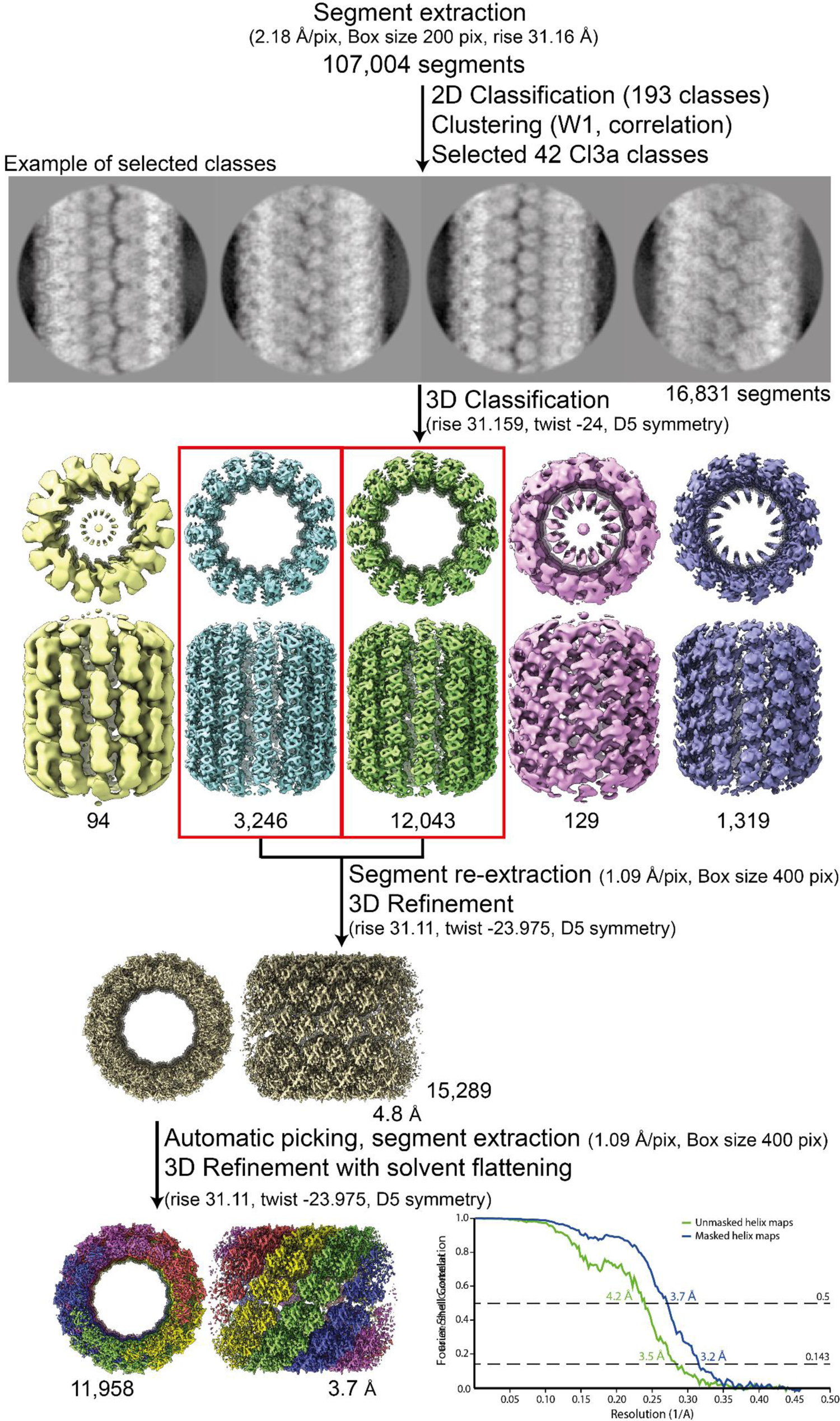
Workflow of the Cl3a relaxed helix reconstruction process and fitting. Segment extraction was performed with a box size of 400 pixels (pix) binned twice (box size 200 pix, 2.18 Å/pix) using the rise obtained from the power spectrum indexation (see methods). From 2D classifications with 200 classes, 42 classes were selected for Cl3a after clustering (see methods and Fig. S5). First, 3D classification was carried out using the 42 2D classes, with D5 symmetry, rise and twist from the power spectrum indexation and a featureless cylinder of 340 Å as reference. 3D-refinement of the two 3D-classes (red boxes) was achieved using one low pass filtered 3D class as reference on the unbinned segments. The resulting map was used as model for automatic picking and segment extraction. Final 3D-refinement was performed on unbinned segments using as initial reference a featureless cylinder with a last step of solvent flattening.

**Fig. S8:**
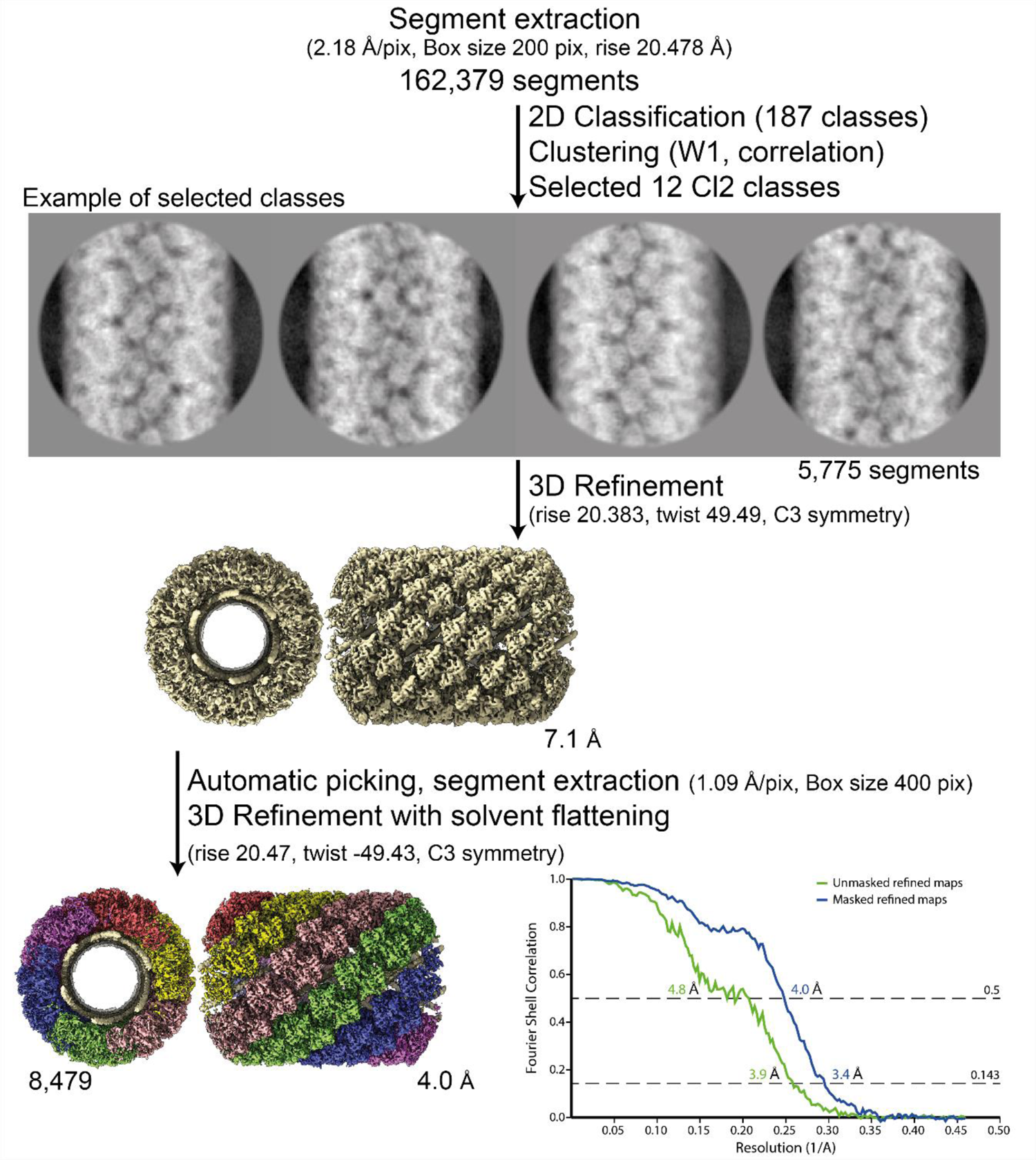
Workflow of the Cl2 helix reconstruction process and fitting. Segment extraction was performed with a box size of 400 pixels (pix) binned twice (box size 200 pix, 2.18 Å/pix) using the rise obtained from the power spectrum indexation (see methods). From 2D classifications with 200 classes, 12 classes were selected for Cl2 based on correlation (see methods and Fig. S5). First, 3D-refinements combining 12 2D classes with solvent flattening, were performed with the C3 symmetry, rise and twist from the power spectrum indexation and a 330 Å featureless cylinder as reference. The resulting map was used as model for automatic picking and segment extraction on the unbinned segments. Final 3D-refinement was performed on unbinned segments using as initial reference a featureless cylinder with a last step of solvent flattening.

**Fig. S9:**
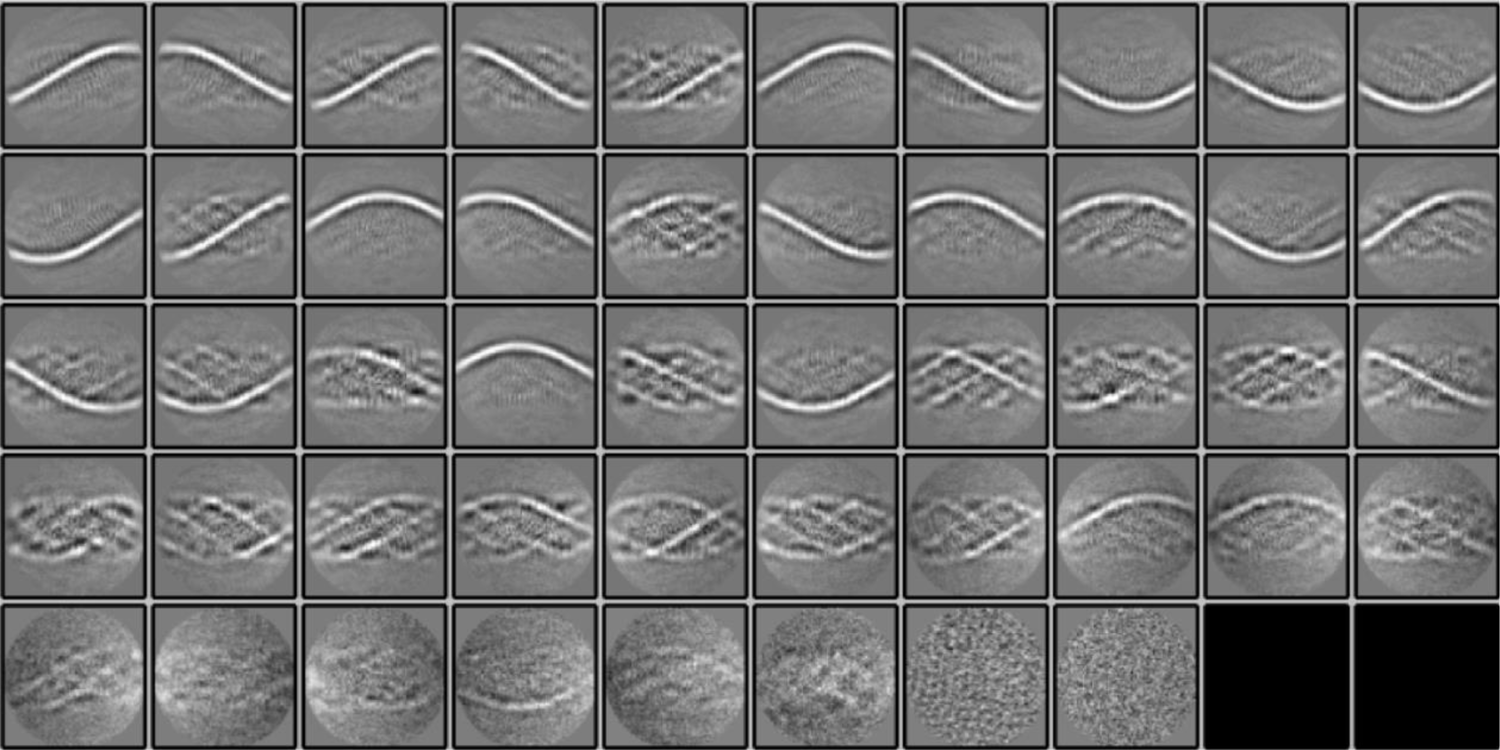
2D Classification of subtracted segments. 50 2D classes (2 empty) were obtained after reference-free 2D classification in Relion 3.1.0 of segments (box size of 400 pixels) of the Cl1a genomic fibers after signal subtraction to keep only the information of the DNA strands inside the fiber. The 2D classes show the enhancement of the signal corresponding to the DNA.

**Fig. S10:**
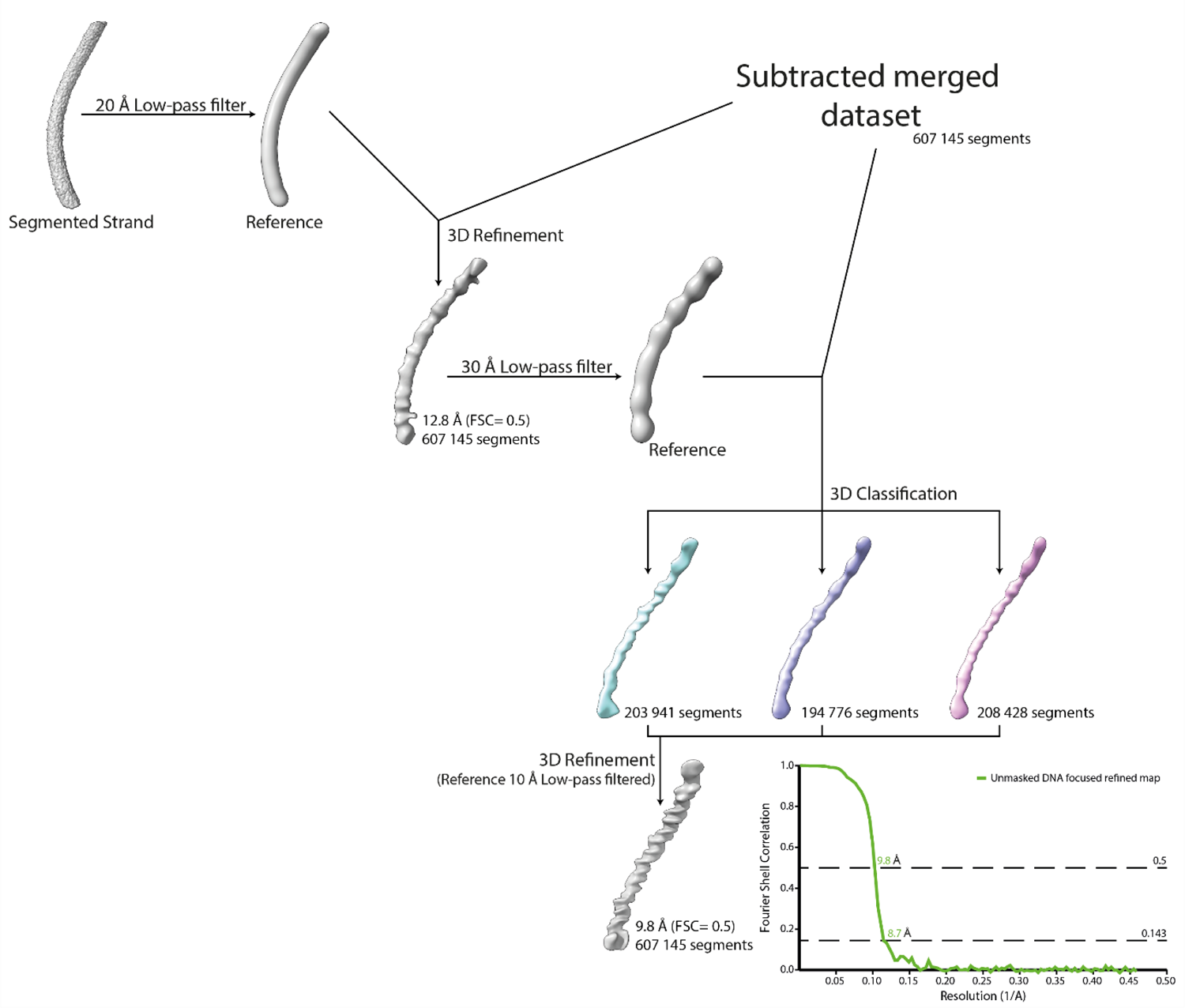
Workflow of the DNA strand focused refinement process. One strand segmented from the 5-start compacted map (Cl1a) was low-pass filtered at 20 Å and used as reference for 3D-refinement of the dataset corresponding to the merged dataset of the individually subtracted 5 strands. The result of the refinement was low-pass filtered at 30 Å and used as reference for 3D-classification. The best 3D class was further refined leading to a focused refined map of the DNA strand.

**Fig. S11:**
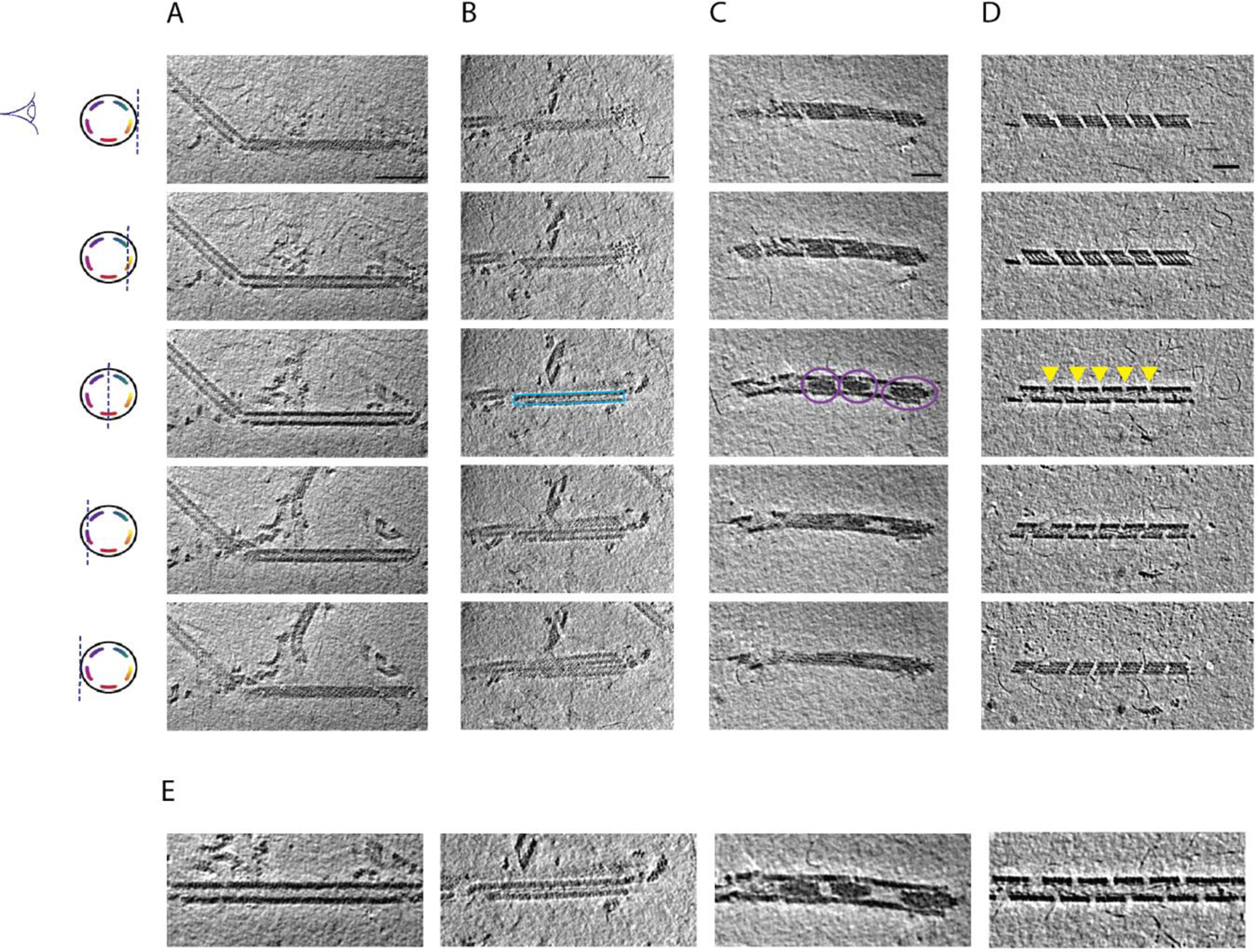
Cryo-ET of Mimivirus genomic fiber. (Upper panel) Slices through tomograms exhibiting different features of compact, unwinding and broken genomic fibers of Mimivirus. (**A**) Slices through a cryo-tomogram focused on a long and broken fiber. Features of the protein shell are visible in tangential slices (top and bottom), while filamentous densities corresponding to DNA strands become visible when slicing deeper inside the fiber. Scale bar, 100 nm. (**B**) A straight filament running along the fiber symmetry axis has been observed in several cases (blue rectangle), which may correspond to a single or a bundle of DNA strands dissociated from the protein shell. (**C**) Large amorphous electron dense structures have been observed inside the lumen of the fiber (highlighted with purple circles). (**D**) DNA strands are often observed emanating from breaks along the genomic fiber, partially disassembling (indicated with yellow arrowheads). (**E**) Detail of the central slices through the genomic fibers showed in the third depicted slice in **A-D**. Thickness of the slices is 1.1 nm. Distance between the tomographic slices (from top to bottom) is 4.4 nm between the first and second and the fourth and fifth, and 6.6 nm between the third and second or fourth. The scheme on the left represents a cross-section of a cartoon representation of the 5-start genomic fiber depicted as a cylinder containing 5 strands of coloured DNA internally lining the helical protein shell. It also indicates the plane through which the genomic fiber is viewed as 2D slices extracted from the tomograms in Tomo. S1 (**A**), Tomo. S2 (**B**), Tomo. S3 (**C**), and Tomo. S4 (**D**). Scale bars (**B**, **C**, **D**), 50 nm.

**Fig. S12:**
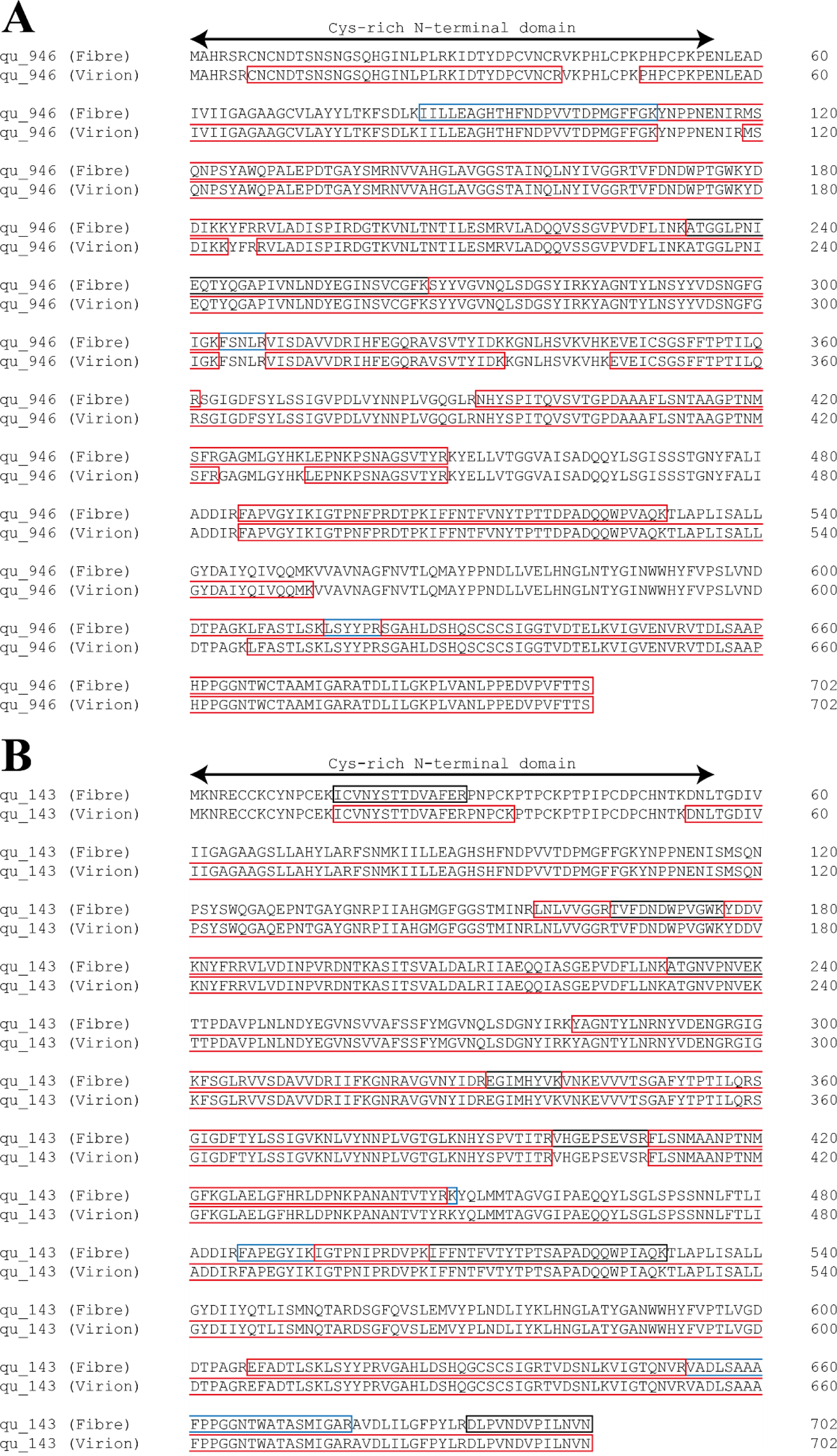
Comparison of sequence coverages for qu_946 (A) and qu_143 (B) obtained by MS-based proteomic analysis of genomic fibers and total virions. Identified peptides are highlighted in red, blue and grey when identified respectively in three, two and one replicate of the purified genomic fiber.

**Table S1:**
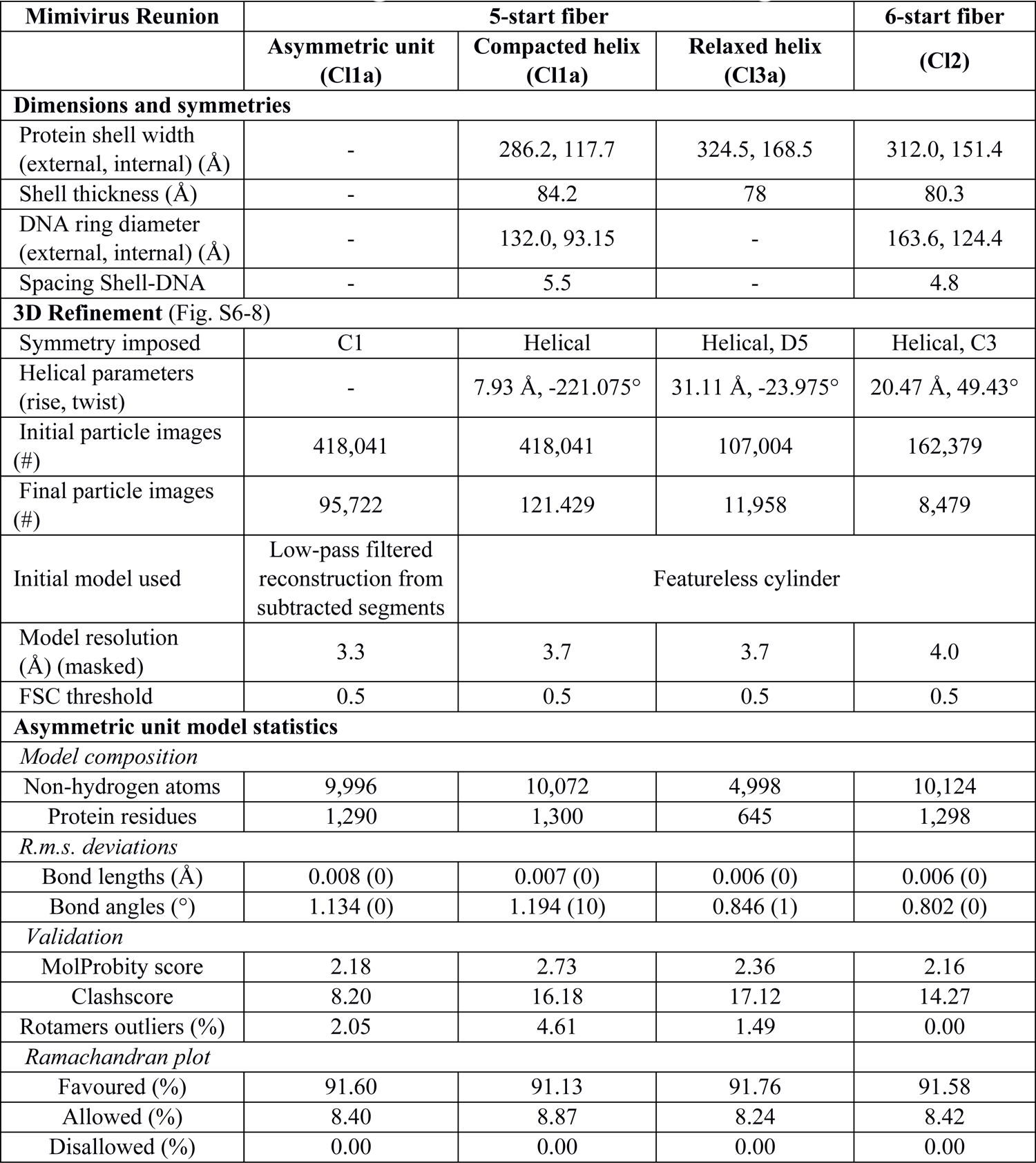
Mimivirus Reunion genomic fibers data statistics (***33***)

**Table S2:**
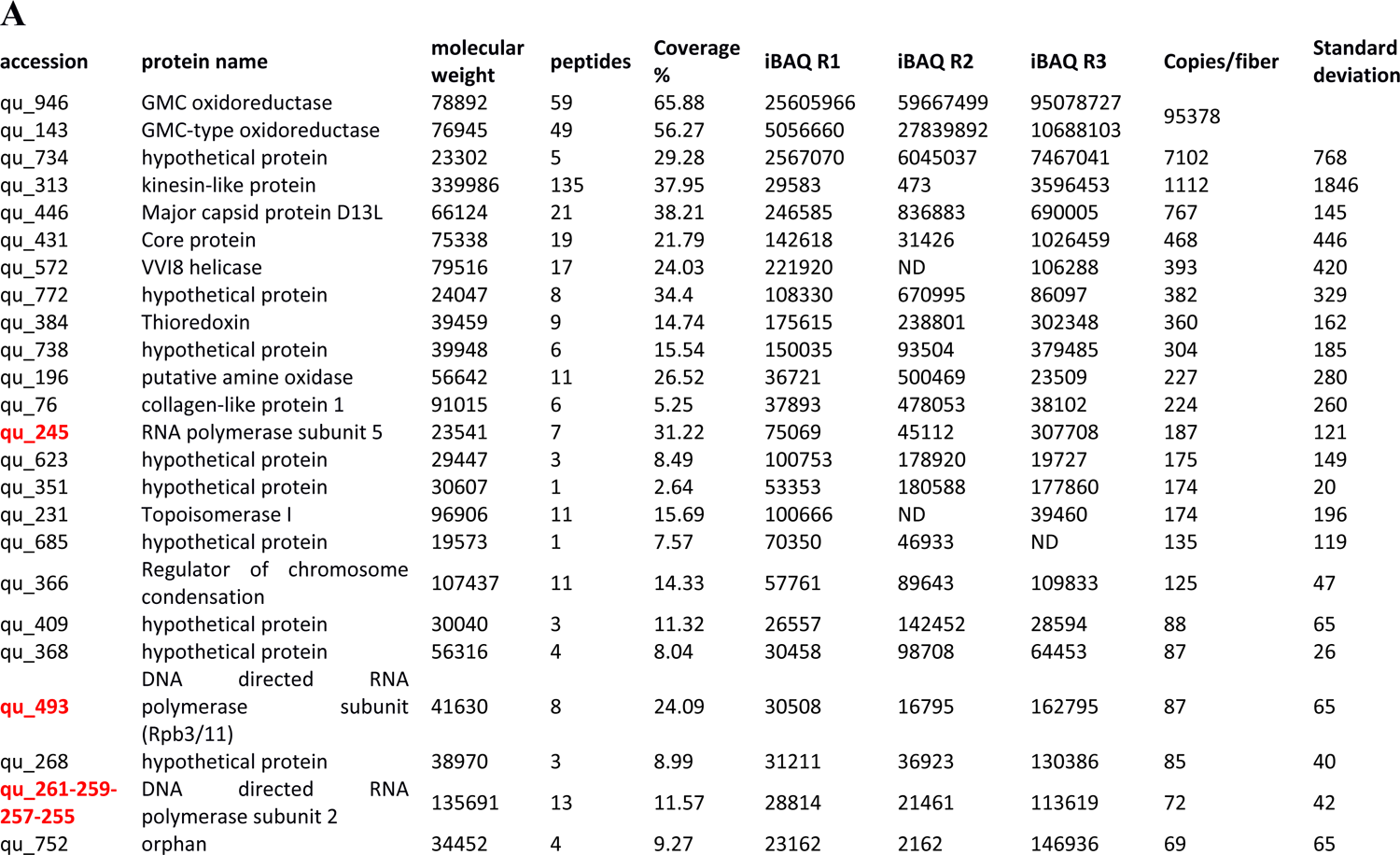
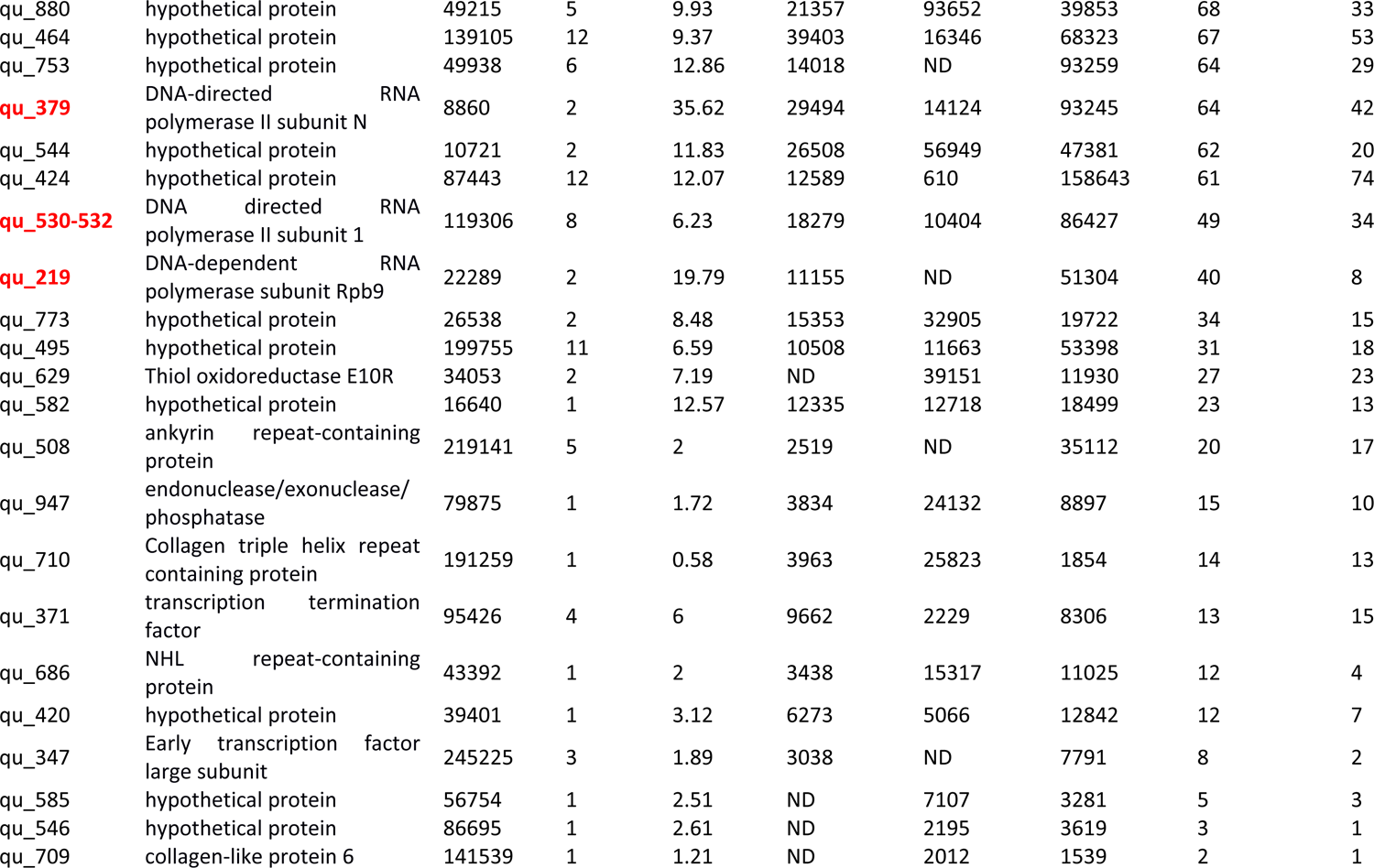
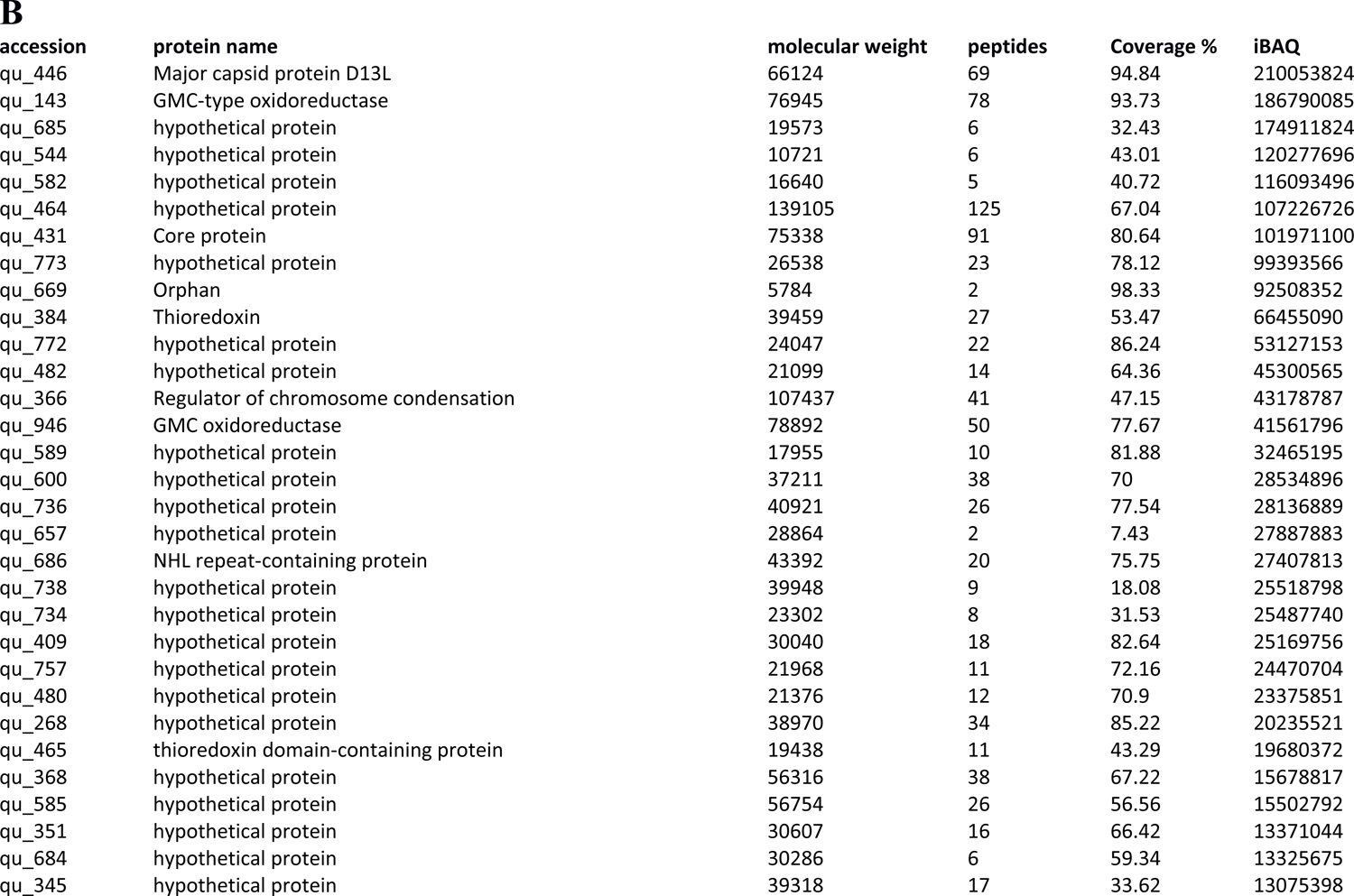
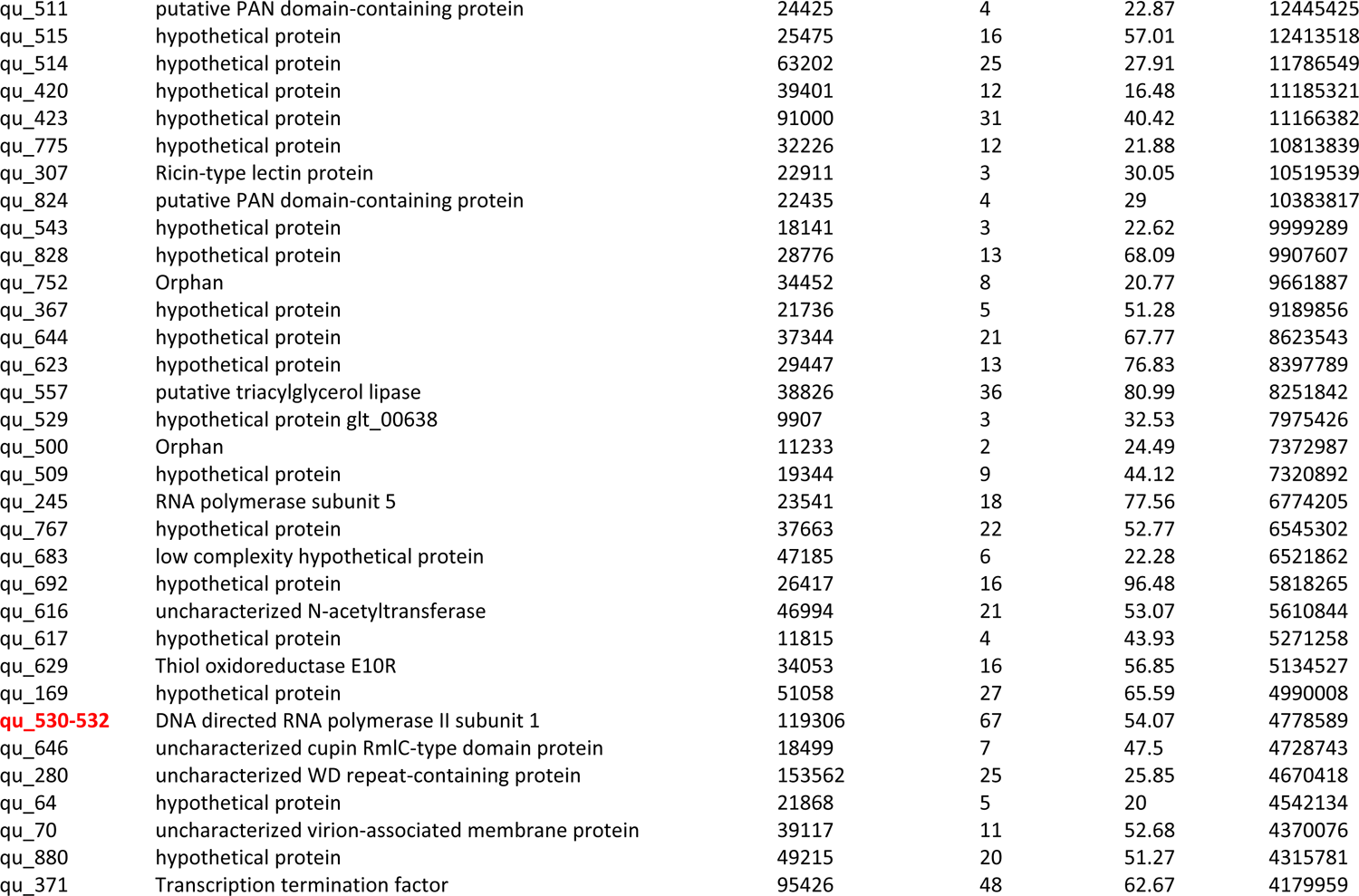
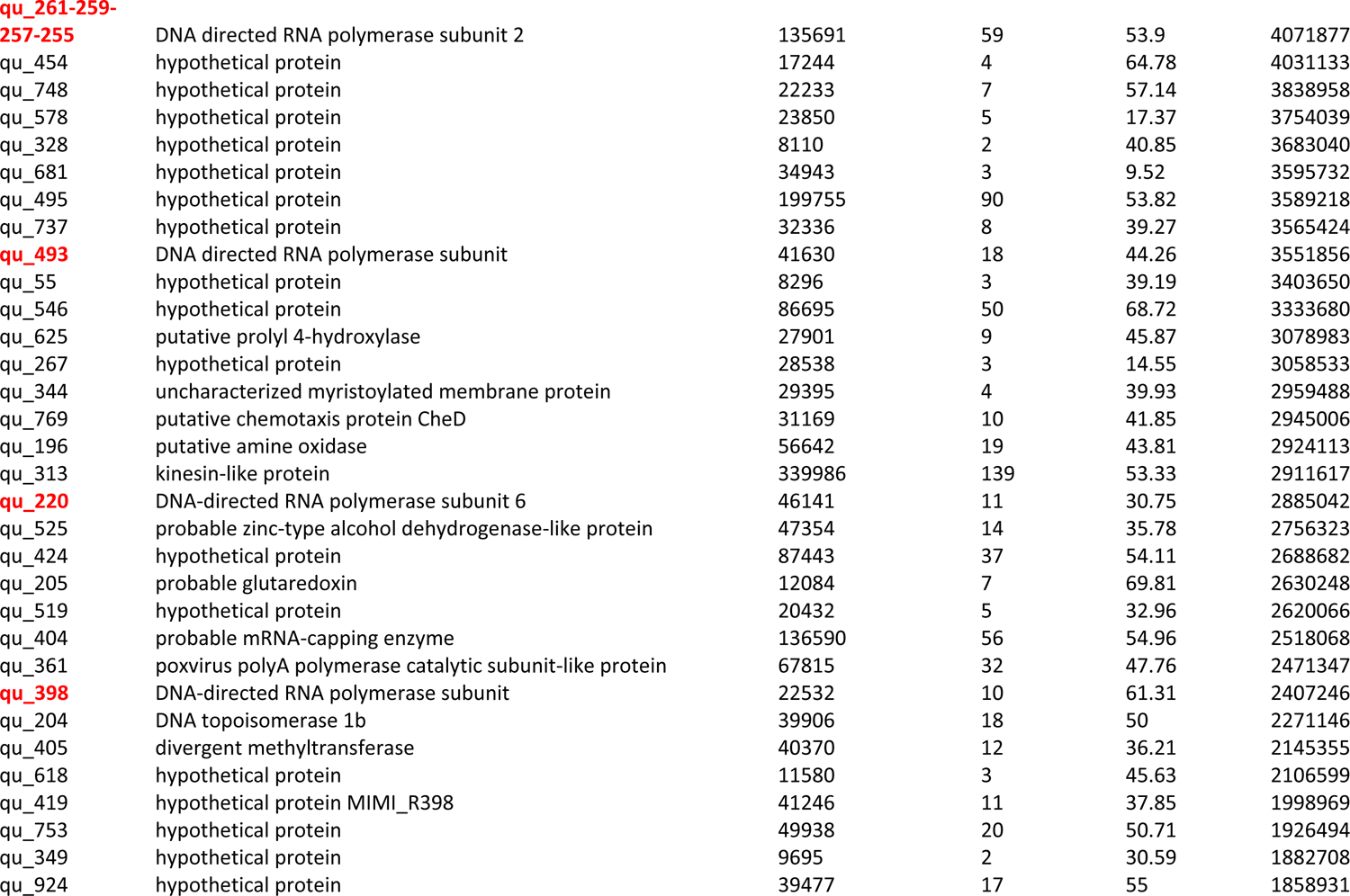
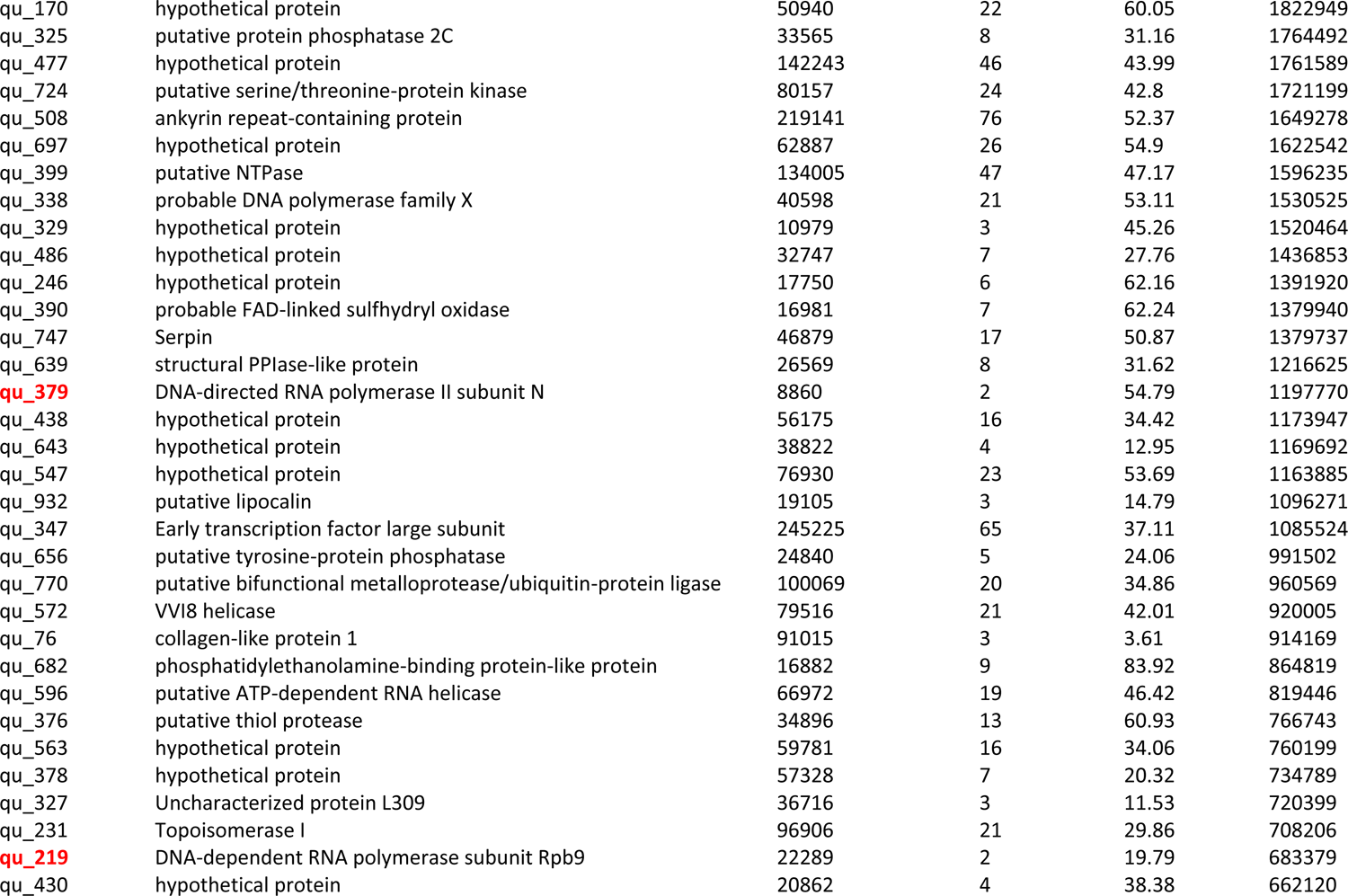
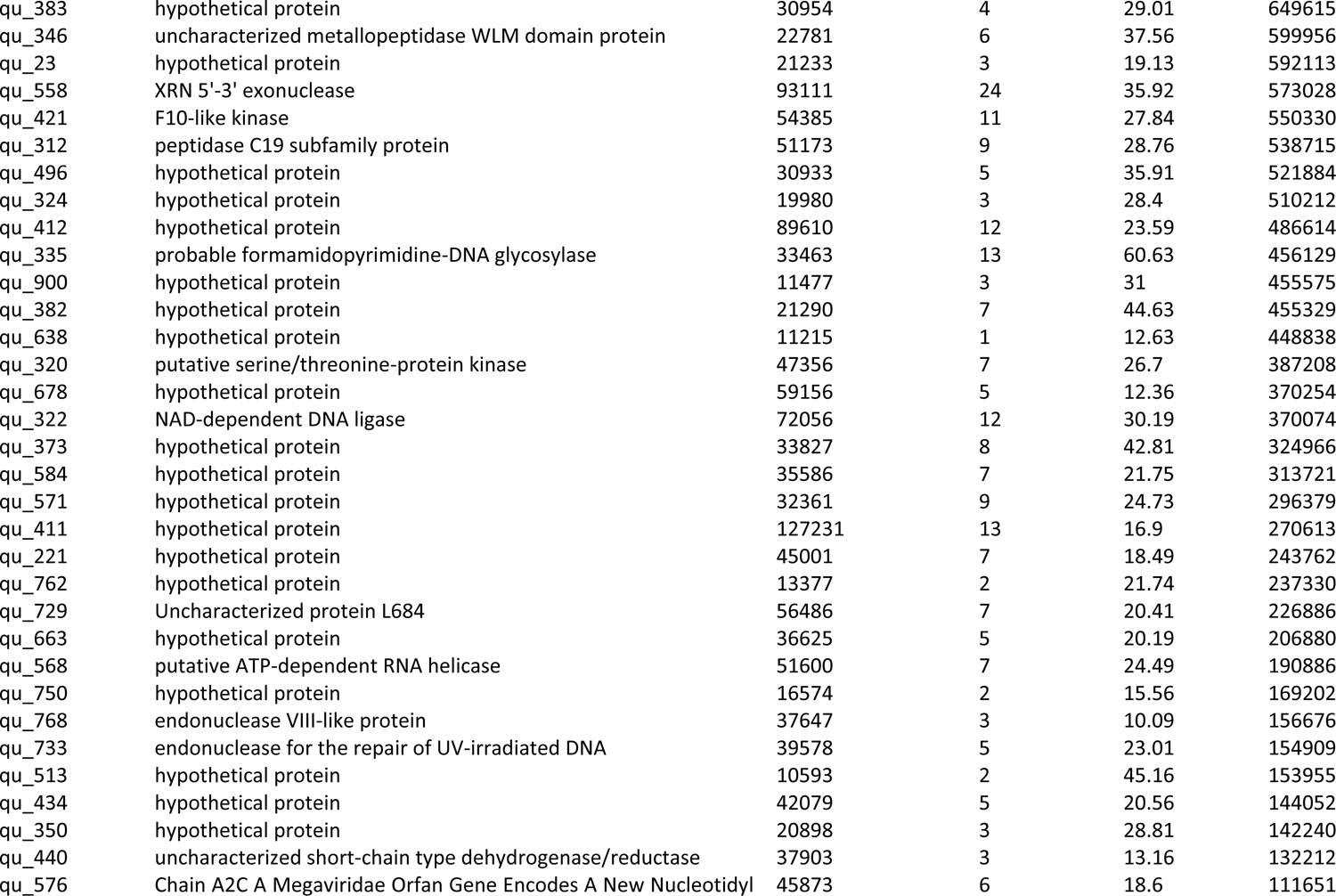
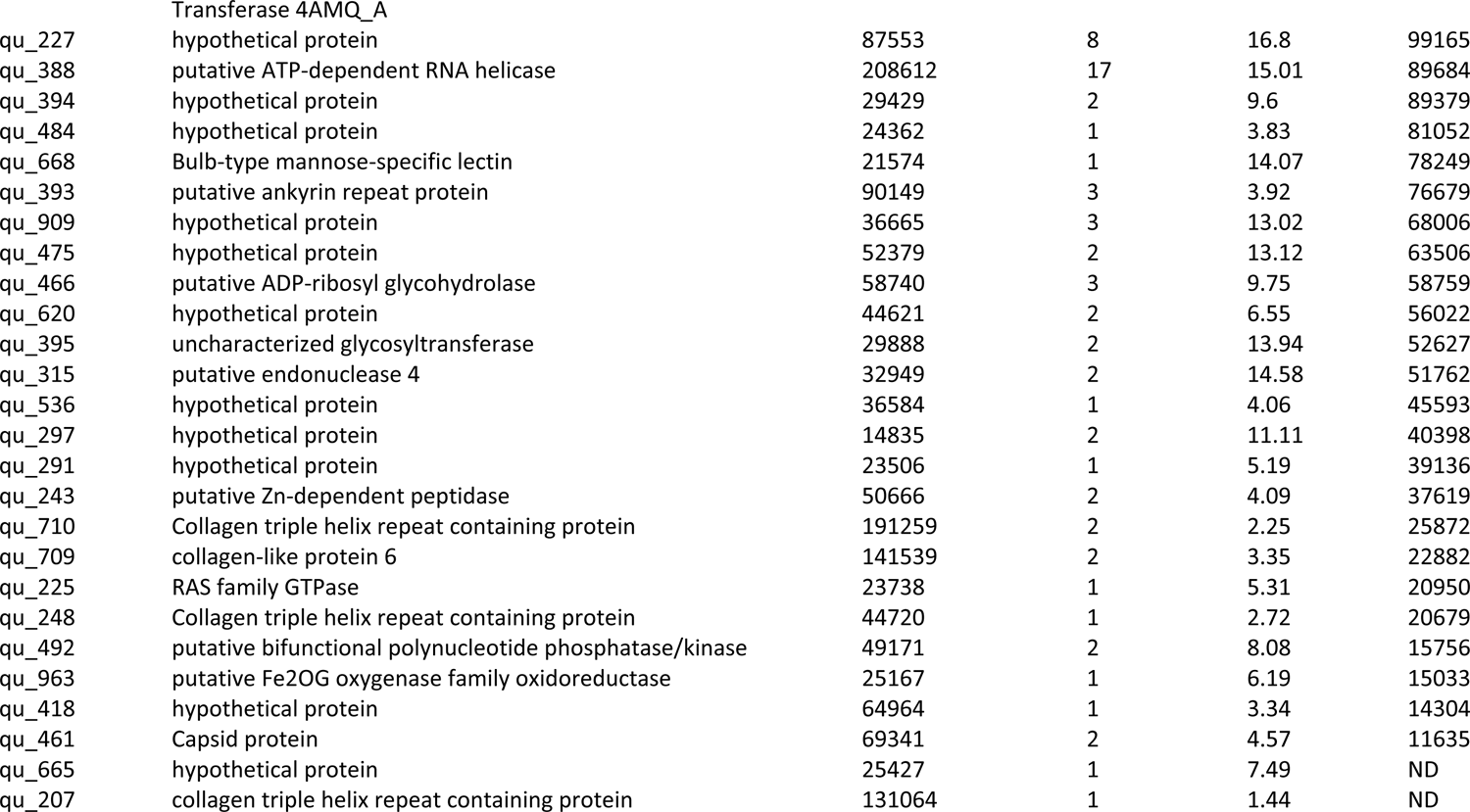
**Mass spectrometry-based proteomic analysis** of A] three independent preparations of Mimivirus genomic fiber and B] one sample of purified Mimivirus virions. (ND: Not detected). RNA polymerase subunits are marked in red.

**Table S3:**
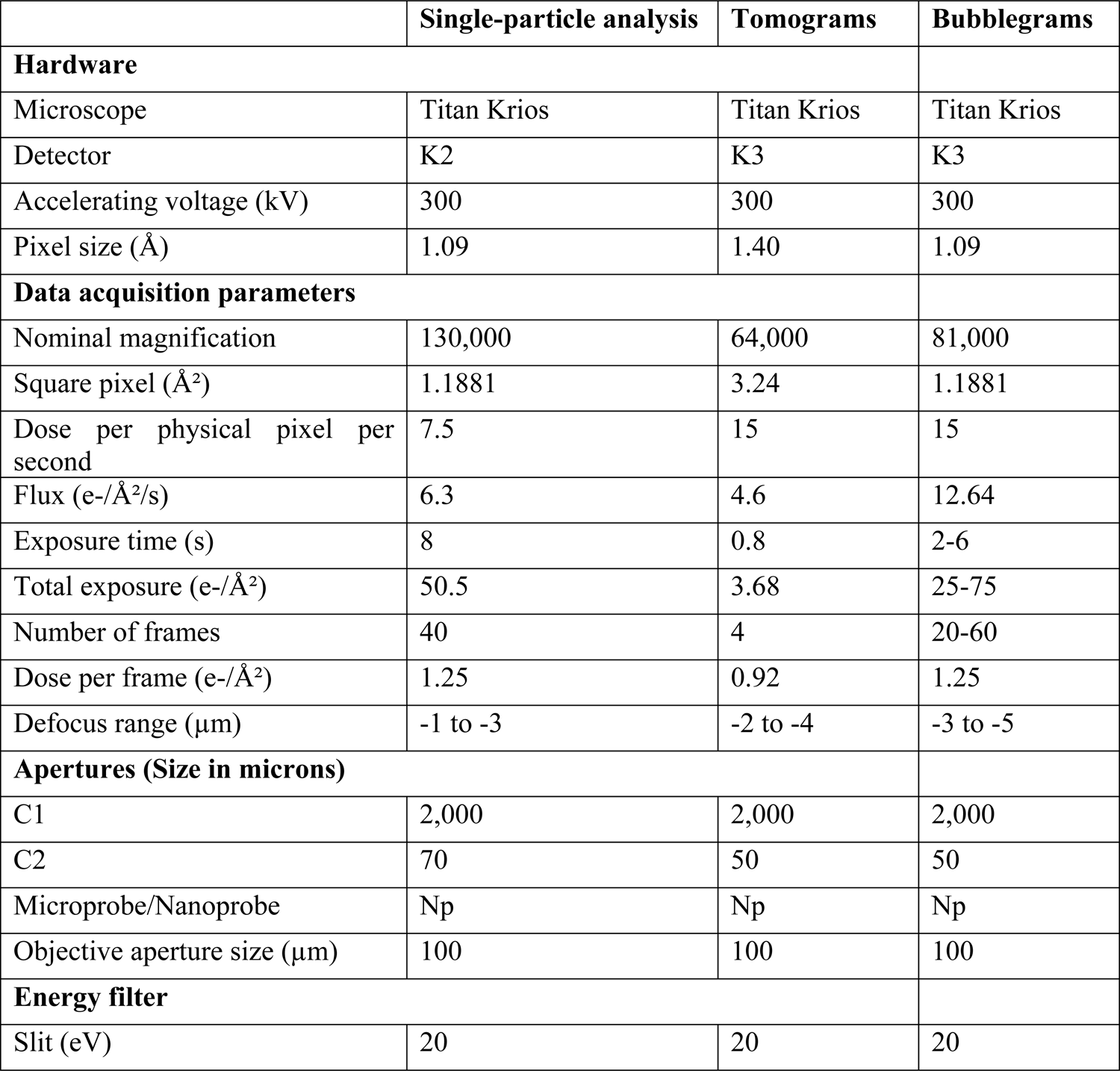
Data acquisition parameters for Cryo-EM

**Video S1:**
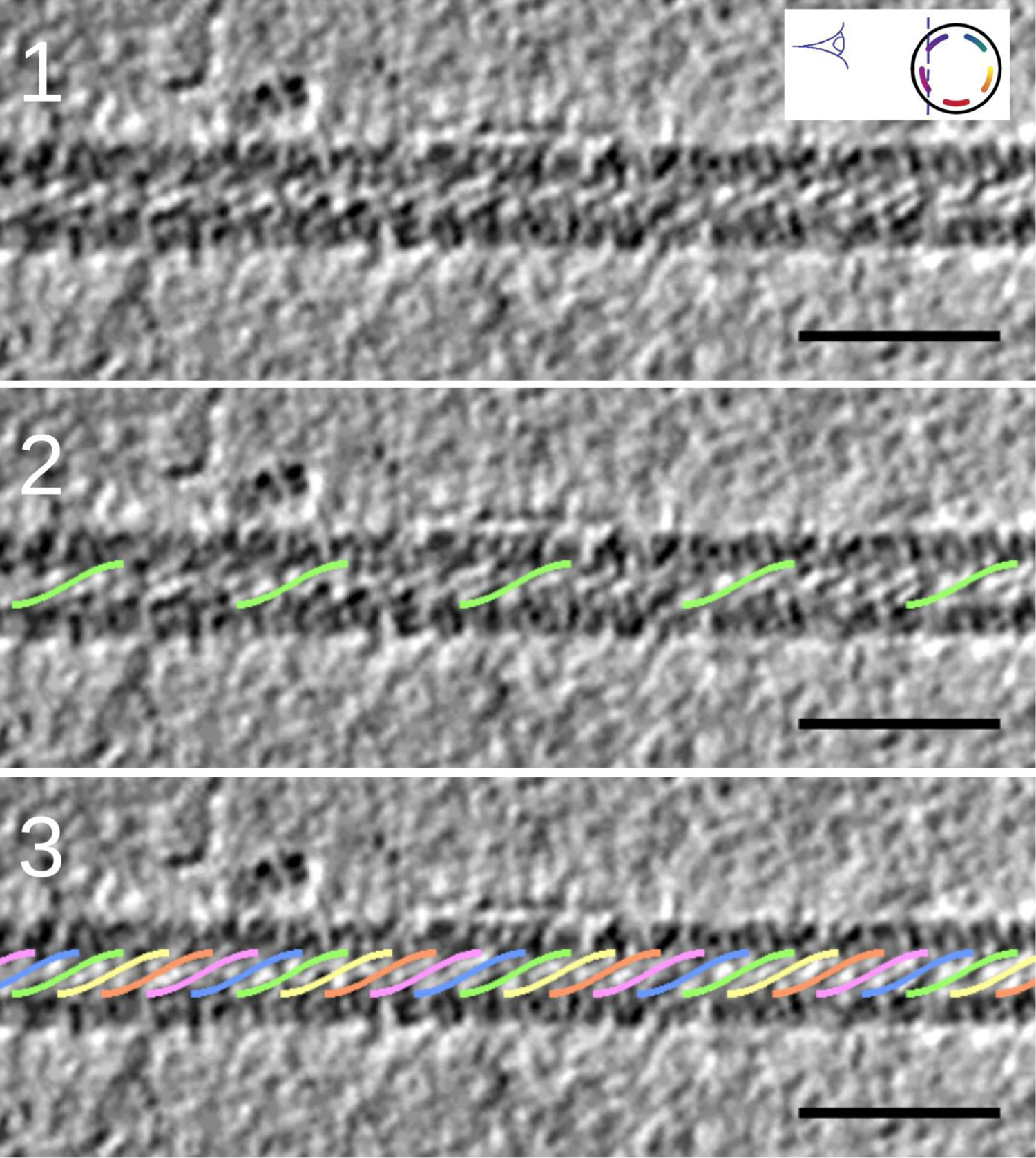
Superimposition of the DNA model on a tomogram slice. 1] one slice of the tomogram where helical strings can be recognized, 2] One strand of the DNA model superimposed on one helical string, 3] 5-start DNA model superimposed on the strings. Scale bar 50 nm.

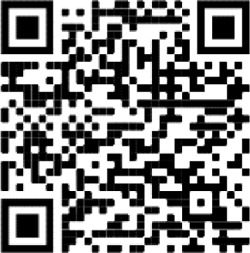

**Video S2:**
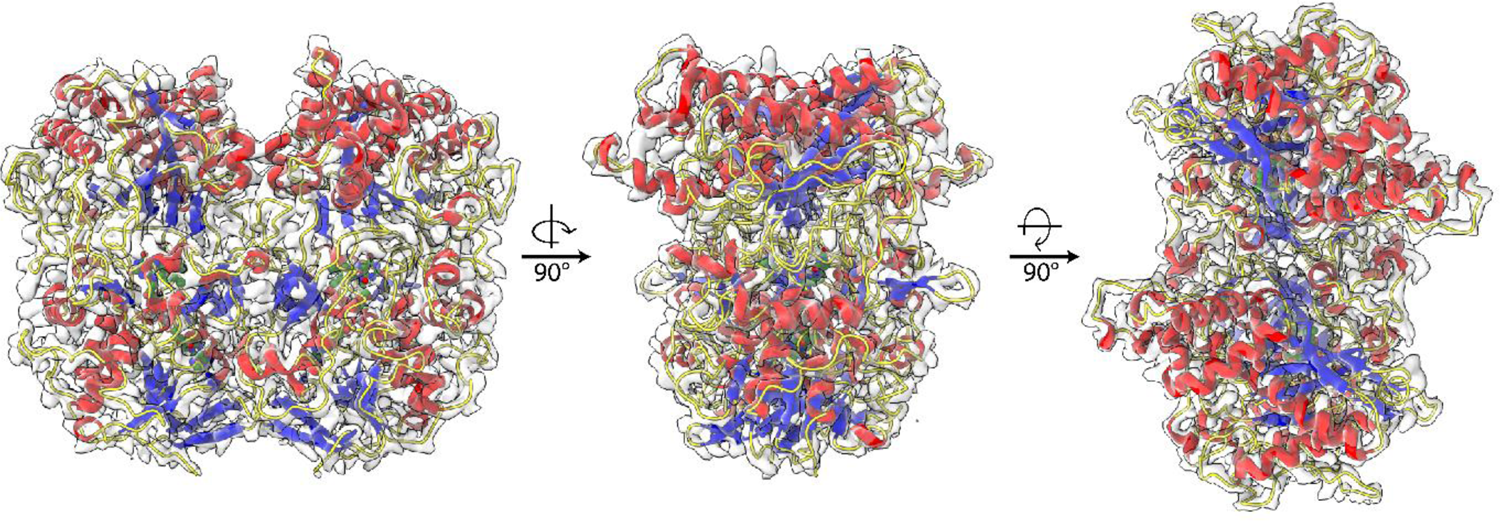
Illustration of the fit of the GMC-oxidoreductase into the asymmetric unit focused refined Cl1a map. View of the overall fit of the GMC-oxidoreductase qu_146 refined model into the Cl1a map with a zoom on the FAD ligand into the map. The video has been made in Chimera 1.14.

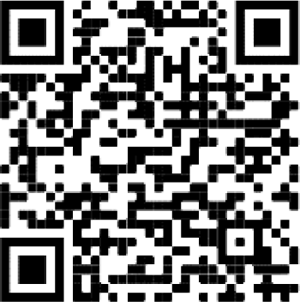

**Video S3:**
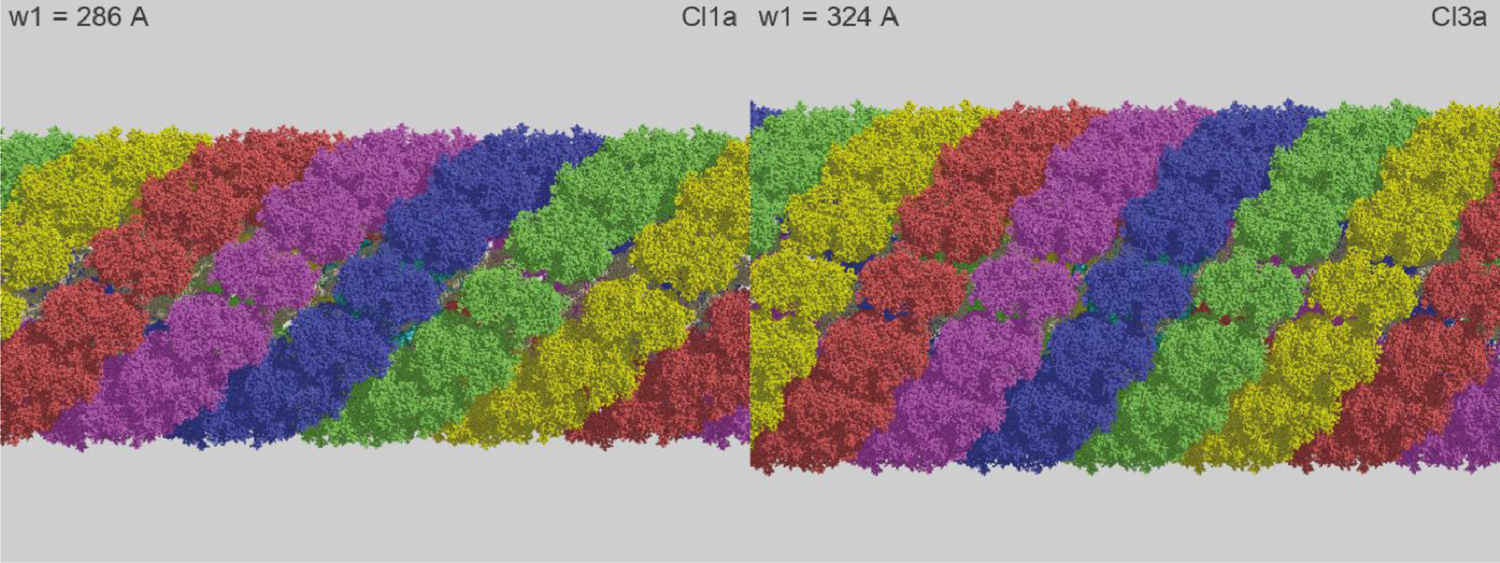
Illustration of the relaxation of the genomic fiber: The relaxation of the helical structure from Cl1a to Cl3a induces a loss of the DNA strands prior to the loss of one start of the protein shell after another once the fully relaxed structure appears as a ribbon. The RNA polymerase is shown in cyan. This video is based on two helical structures described in this work, at relaxing states corresponding to Cl1a and Cl3a, while all intermediate states were obtained by linear interpolation of the helical parameters.

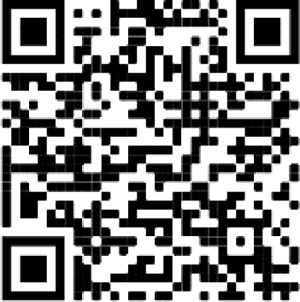

